# Systemic genome-epigenome analysis captures the lineage specificity and functional significance for *MYB* associated super-enhancer in gastrointestinal adenocarcinoma

**DOI:** 10.1101/2024.11.11.622904

**Authors:** Fuyuan Li, Shangzi Wang, Lian Chen, Ning Jiang, Xingdong Chen, Jin Li

## Abstract

The gastrointestinal adenocarcinoma is a major cancer type for the digestive system, ranking as the top cause of cancer-related deaths worldwide. In contrast to the large body of studies on the protein-coding regions’ mutations, the knowledge about the landscape of its non-coding regulatory elements is still insufficient. Combining the analysis of active enhancers profile and genomic structural variation, we discovered and validated a lineage-specific super-enhancer for *MYB* in gastrointestinal adenocarcinoma. This super-enhancer is constituted by a predominant enhancer e4 and multiple facilitator enhancers, whose transcriptional activity is controlled by the direct binding of HNF4A and MYB itself. Suppression of the super-enhancer downregulated the expression of *MYB*, inhibited the downstream Notch signaling and prevented the development of gastrointestinal adenocarcinoma *in vitro* and *in vivo*. Our study revealed a non-coding variation-based mechanism to affect *MYB* expression in a lineage-specific manner, which provided an inspiring insight into the carcinogenic mechanism and therapeutic strategies for gastrointestinal adenocarcinoma.

## Introduction

The overexpression of oncogenes in patients contributed to the development of cancer, some of which became therapeutic targets and inspired the pharmaceutic industry to develop better therapies for cancer^1–3^. However, the majority of these studies focused on the structural variations or gain-of-function mutations in the protein-coding regions of human genome, which only accounts for a part of cancer associated genetic variations^4–7^. It has been validated that the variations of non-coding region in human genome have significant impacts on health^8–11^. Therefore, it is reasonable to hypothesize that the non-coding variants also play an important role in the expression program of the oncogenes as well as the progress of cancer. By nature, the transcriptional regulation by the non-coding regions is highly lineage-specific. Thus, research is needed to better understand the role of non-coding regions on cancer development and progression and uncover new therapeutic targets for individual cancer type. For instance, gastrointestinal adenocarcinoma is a major cancer types for the digestive system, ranking as the top cause of cancer-related deaths worldwide^12^. However, knowledge about its lineage-specific variations in the non-coding regions is still lacking.

Dysregulation of transcriptional programs can be mediated by epigenetic alterations targeting the non-coding regulatory elements like enhancers and super-enhancers (SEs). In the context of cancer research, SE has been described as a cluster of enhancers in close proximity, which are bound by H3K27ac, transcription factors or mediator complex to drive the high expression of lineage-specific oncogenes^13^. It has been demonstrated that the recurrent alterations of SEs play an important role in tumorigenesis and malignant phenotypes by interacting with the corresponding promoters via the three-dimensional (3D) structure of genome^14–17^. For example, SE amplifications increase expression of oncogenes including *MYC*, *SOX2*, *USP12*, *KLF5*, *ZFP36L2* and control the progress of various kinds of tumors including endometrioma and gastrointestinal tumors^14–17^. Considering the diversity of different types of cancer, remarkable efforts are demanded to understand the transcriptional regulation of oncogenes by the SEs in gastrointestinal adenocarcinoma, especially colorectal cancer (CRC) and gastric cancer.

*MYB* is a protein-coding gene containing a highly conserved DNA-binding domain^18^. As a transcription factor, it controls the expression of several oncogenes including *MYC*, *BIRC5* and *IGF1R*^19^. The overexpression of *MYB* is widely observed in cancer cells and the oncogenic activity of *MYB* has been confirmed in leukemia, breast cancer, prostate cancer, et al^19, 20^. Most of the known mechanisms for the overexpression of *MYB* are related to gene duplication, gene translocation or intragenic mutation^19, 21–24^. But the regulation of *MYB* expression by non-coding variation in human genome has not been investigated.

In the current study, we identified a lineage-specific SE for gastrointestinal adenocarcinoma, which robustly enhances the expression of *MYB* and controls the development of tumor. A single predominant enhancer e4 of this SE was discovered, whose activities were controlled by the direct binding of HNF4A/MYB and affected by the interaction with other enhancers in the SE. Suppressing the activity of e4 downregulated the expression of *MYB* and the downstream Notch signaling, reduced the cell viability of CRC and gastric cancer cells and prevented the growth of tumor xenograft. Our study revealed a non-coding variation-based mechanism to affect *MYB* expression in a lineage-specific manner, which provided an inspiring insight into the carcinogenic mechanism and therapeutic strategies for gastrointestinal adenocarcinoma.

## Results

### A lineage-specific super-enhancer near the *MYB* gene is duplicated in gastrointestinal adenocarcinoma

SEs are acquired by cancer cells at key oncogenes^13^. We investigated the landscape of active enhancers both in primary tumors and cell lines for colon and rectum adenocarcinoma (COREAD) and stomach adenocarcinoma (STAD) by using the data of chromatin immunoprecipitation-sequencing (ChIP-seq) based on H3K27ac, a marker for activate promoters and enhancers, from various dataset deposited at Gene Expression Omnibus (GEO)^15, 25–28^. We identified 7897 and 4113 SEs in cancer cell lines and tumor samples, respectively (**Fig. 1A**). Considering the shared lineage between colon and stomach, we ought to focus on the SEs that shared by COREAD and STAD. Cell lines and tumors shared 82 and 185 SEs assigned genes, respectively, adjacent 205 unique protein-coding genes (**Fig. 1B and Supplementary Table 1**) including 34 oncogenes (**Fig. 1C**). Gene Ontology (GO)^29^ and Kyoto Encyclopedia of Genes and Genomes (KEGG)^30^ pathway analyses revealed that these genes were predominantly associated with transcriptional regulation and cancer (**Fig. 1D and Supplementary Fig. 1A**). Furthermore, Reactome pathway analysis showed that these genes were involved in canonical cancer-related signaling pathways, such as Notch and TGF-β signaling. (**Supplementary Fig. 1B**). These findings suggested that the common SEs for may play an important role for the development of gastrointestinal adenocarcinoma.

**Figure 1.**
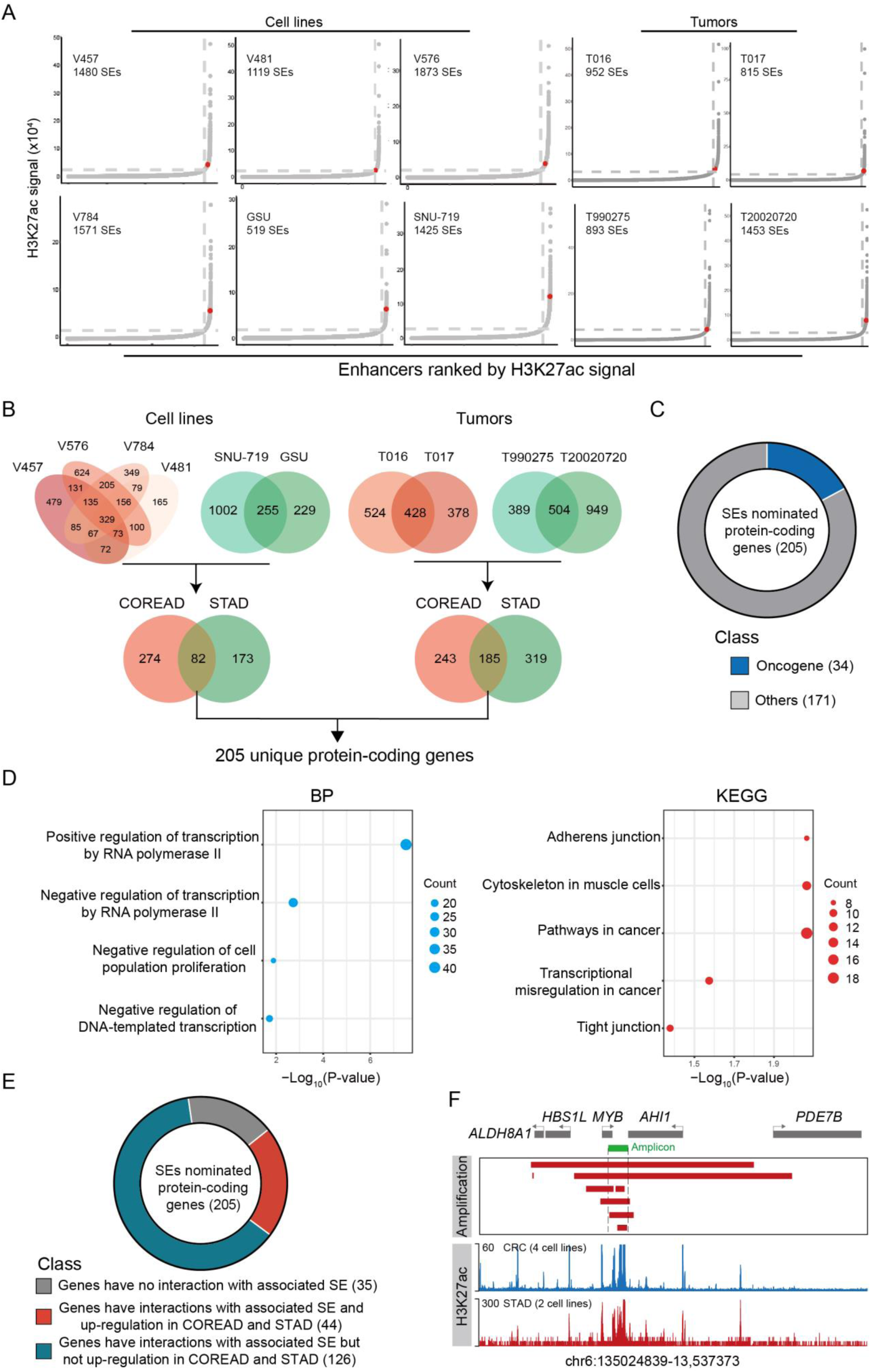
A super-enhancer near *MYB* locus is duplicated in gastrointestinal adenocarcinoma. **A.** Hockey stick plots for super-enhancer profiles of COREAD and STAD in cancer cell lines. The ranking was calculated by the H3K27ac ChIP-seq signal enrichment. The super-enhancers associated with *MYB* were highlighted in red; **B.** Venn diagram showing the shared super-enhancers of various gastrointestinal cancer cell lines or tumor samples respectively; **C.** Pie chart showing oncogene classification of 205 SEs nominated protein-coding genes; **D.** The enrichment and functional analysis of 205 SEs nominated protein-coding genes. BP: biological process. KEGG: Kyoto Encyclopedia of Genes and Genomes; **E.** Pie chart showing categorization of 205 SEs nominated protein-coding genes based on enhancer-promoter interaction and expression; **F.** The *MYB* amplicons across gastrointestinal adenocarcinoma samples and merged enhancer profile from cell lines at the *MYB*-associated loci in gastrointestinal adenocarcinoma.

Among the genes associated to common SEs, 44 genes had physical interactions with SEs based on H3K27ac HiChIP analysis and were upregulated in COREAD and STAD based on RNA-seq analysis (**Fig. 1E and Supplementary Fig. 1C**). Among them, 11 genes were previously annotated oncogenes that have physical interactions with their SEs and upregulated both in COREAD and STAD (**Supplementary Fig. 2A**). As a common event of genomic structural variation in cancer cells, duplication often drives abnormally high expression of oncogenes. In a higher-resolution whole-genome sequencing dataset from the Pan-Cancer Atlas of Whole Genomes (PCAWG) Project^31^, we observed a recurrent amplicon in the noncoding regions near *MYB* in gastrointestinal adenocarcinoma, an oncogene widely expressed in acute myeloid leukemia (AML)^19, 20^. Especially, *MYB* was a gene with SE present in both tumor samples and cell lines (**Supplementary Fig. 2B**). The SE (*MYB*-SE) inside the duplication hotspot near the *MYB* was presented at **Fig. 2F** and **Supplementary Fig. 2C**.

**Figure 2.**
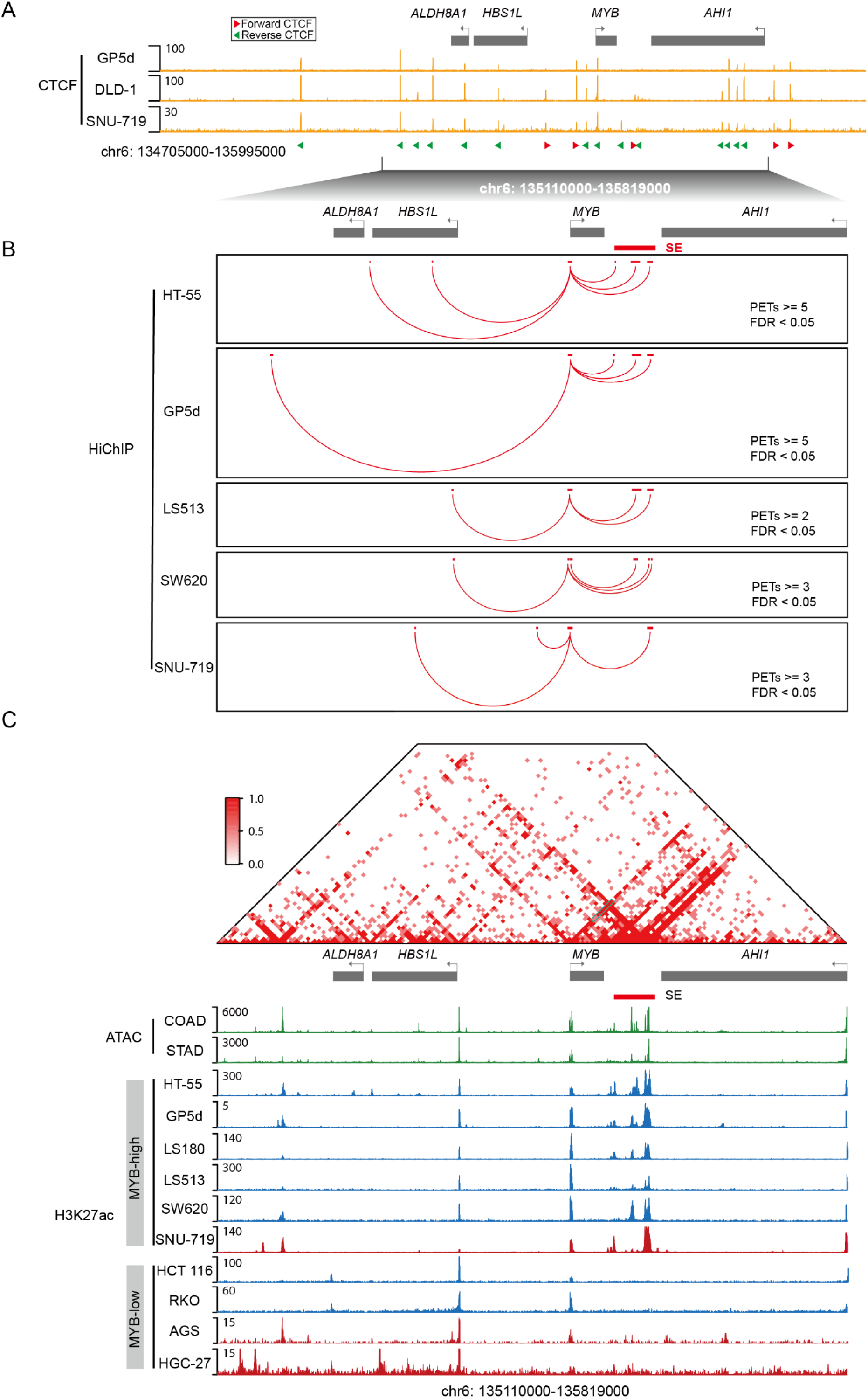
Long-range interaction between the super-enhancer and promoter of MYB in gastrointestinal adenocarcinoma. **A.** The CTCF ChIP-seq tracks of multiple gastrointestinal cancer cell lines were presented. The red and green arrows represented the orientation of the CTCF motifs; **B.** The chromatin loops based on H3K27ac HiChIP data connected the *MYB* promoter to multiple distal elements with high-confidence in various gastrointestinal cancer cell lines; **C.** The demonstration of the epigenetic features for the *MYB* associated loci. Top: The chromatin interaction frequency heatmap in the gastrointestinal cancer cell line HT-55 based on H3K27ac HiChIP data. Turquoise box showed the interactions between *MYB* promoter and *MYB-*SE. Middle: The ATAC - seq tracks of COREAD and STAD samples. Bottom: The H3K27ac ChIP - seq tracks of 10 gastrointestinal cancer cell lines. Blue for COREAD tracks and red red STAD tracks.

It has been reported that the activity of SEs is cell-type and tumor-type specific^13, 32^, so we examined the lineage specificity of the candidate *MYB*-SE in other types of cancer. Although the genomic region of the candidate *MYB*-SE was also duplicated in esophageal adenocarcinoma (ESAD), a cancer type with close lineage association but remarkable biological differences with COREAD and STAD, the H3K27ac ChIP-seq analysis revealed that it did not constitute a SE (**Supplementary Fig. 2D**). In addition, though *MYB* is considered as a cancer oncogene in AML with a potential super-enhancer^33^, the candidate *MYB*-SE was not presented in AML and many other types of cancer (**Supplementary Fig. 2D**). We also tested the existence of *MYB*-SE in other species with a published dataset from mouse^34^. In comparison to *Myc*, the low H3K27ac ChIP-seq signal (**Supplementary Fig. 2E**) and low expression of *Myb* was identified in mouse CRC cell (**Supplementary Fig. 2F**), indicating *MYB*-SE is likely to be specific for human.

May the existence of the candidate *MYB*-SE have any impacts on the expression of *MYB* in gastrointestinal adenocarcinoma? Based on RNA-seq data of normal samples from the Genotype-Tissue Expression project (GTEx)^35, 36^ and tumor samples from the Cancer Genome Atlas (TCGA)^37^, the expression of *MYB* in COAD, READ and STAD tumors was higher than normal tissue (**Supplementary Fig. 2G**). Similar results were obtained from the expression profiles of paired samples from TCAG (**Supplementary Fig. 2H**). Using cell lines data from CCLE^38^, we found that the expression of *MYB* only presented a weak correlation with the copy number of its coding region (**Supplementary Fig. 2I**), indicating the copy number may not be a major driver for the expression of *MYB*. Thus, we hypothesized that the overexpression of *MYB* in COREAD and STAD was caused by the presences of the duplicated candidate *MYB*-SE.

### 3D genomics analysis identified *MYB* candidate functional enhancers in gastrointestinal adenocarcinomas

We next sought to explore how the candidate *MYB*-SE regulated the expression of *MYB* gene in gastrointestinal cancer. It is widely accepted that the three-dimensional architectures of chromatin were important for the transcriptional regulation. Therefore, we firstly investigated the interaction between *MYB* promoter and candidate *MYB*-SE with a published dataset of ChIP-seq based on the CCCTC-binding factor (CTCF), which plays an indispensable role in mediating the formation of topologically associating domains (TADs) and chromatin loops^39–42^. Interestingly, *MYB* promoter and candidate *MYB*-SE resided within the same adjacent insulated neighborhoods that was defined by mutually convergent CTCFs (**Fig. 2A**), suggesting that *MYB* promoter may interact with candidate *MYB*-SE.

By analyzing the published H3K27ac HiChIP data in representative cell lines (**Supplementary Table 2**)^25, 39, 43–46^, we investigated the physical interactions between *MYB* promoter and the enhancers residing in candidate *MYB*-SE as promoter-enhancer loops. The expression and dependency for *MYB* of the COREAD and STAD cell lines were presented in **Supplementary Fig. 3A**, based on the data from DepMap^47^. The results of HiChIP analysis indicated that the constituent enhancers of candidate *MYB*-SE had highly confident pair-end-tags (false discovery rate (FDR) < 0.05) to link them to *MYB* promoter in *MYB*-high/-dependent cell lines (Fig. 2B). In contrast, these pair-end-tags were absent in *MYB*-low/-independent cell lines (including HCT 116, RKO, AGS cell lines) and normal human primary colonic epithelial cells (**Supplementary Fig. 3B**).

We next analyzed the chromatin landscape with the data from the Assay for Transposase-Accessible Chromatin with high throughput sequencing (ATAC-seq) in patient samples^48^ and ChIP-seq of H3K27ac in COREAD and STAD cancer cell lines with high or low *MYB* expression (**Fig. 2C**). We found that the candidate *MYB*-SE region exhibited a chromatin-accessible state (**Fig. 2C**), and the *MYB*-highly expressed cancer cell lines presented convincing H3K27ac signals in the candidate *MYB*-SE region (**Fig. 2C**). On the contrary, the *MYB*-lowly expressed cancer cell lines showed minimal enrichment of H3K27ac signal in the candidate *MYB*-SE region (**Fig. 2C**). The situation became more complicated when *MYB* dependency was taken into consideration. We identified the presences of candidate *MYB*-SE in *MYB*-high/-dependent and *MYB*-high/-independent but not *MYB*-low/-independent cell lines (**Supplementary Fig. 3C**). Taken together, these results revealed the correlation between the presence of candidate *MYB*-SE and the expression or dependency of *MYB* in gastrointestinal adenocarcinoma.

### *MYB* activation is predominantly driven by an individual enhancer within the *MYB*-SE

We further explored the importance of *MYB* for gastrointestinal adenocarcinoma by investigating the consequence of its genetic depletion or suppression based on the data from cancer cell dependency analysis by the DepMap project^47^, CRISPR-based genetic screen by the Sanger Institute^49^ and shRNA-based screen project by the Broad Institute and Novartis^50^. The *MYB* dependency level identified from the data of DepMap project was tightly correlates with the results from the other two datasets, indicating the robustness of the analysis (**Supplementary Fig.4A**). Importantly, the cancer cell lines with higher expression levels of *MYB* exhibited an even greater dependent on the *MYB* gene for many types of cancer, including COREAD and STAD (**Supplementary Fig. 4B**). These results highlighted the significance of *MYB* gene overexpression for gastrointestinal adenocarcinoma.

To identify the predominant enhancer(s) within *MYB*-SE which may drive *MYB* overexpression, we divided the *MYB*-SE into seven individual enhancers, according to the signals from ATAC-seq and H3K27ac ChIP-seq analysis in this region (**Fig. 3A**). The initial analysis was performed on the data from self-transcribing active regulatory region sequencing (STARR-seq)^42^, a method to directly identify the enhancers in a genome-wide manner. The results revealed that e4 has stronger transcriptional activity compare to the other enhances of the *MYB*-SE in the *MYB*-high and -dependent GP5d cells (**Supplementary Fig. 4C**). We further validated the significance of e4 for *MYB* expression with the luciferase assay in HT-55 and SNU-719 cells (**Fig. 3B**). In contrast, the activity of e4 was not detected in HEK293-FT cells (**Supplementary Fig. 4D**), which is consistent to the discovery of lineage specificity of *MYB*-SE for gastrointestinal adenocarcinomas. To mimic enhancer duplication in the gastrointestinal adenocarcinomas, we generated a plasmid with an additional copy of e4 (**Supplementary Fig. 4E**), which indeed demonstrated a stronger luciferase activity (**Fig. 3C**).

**Figure 3.**
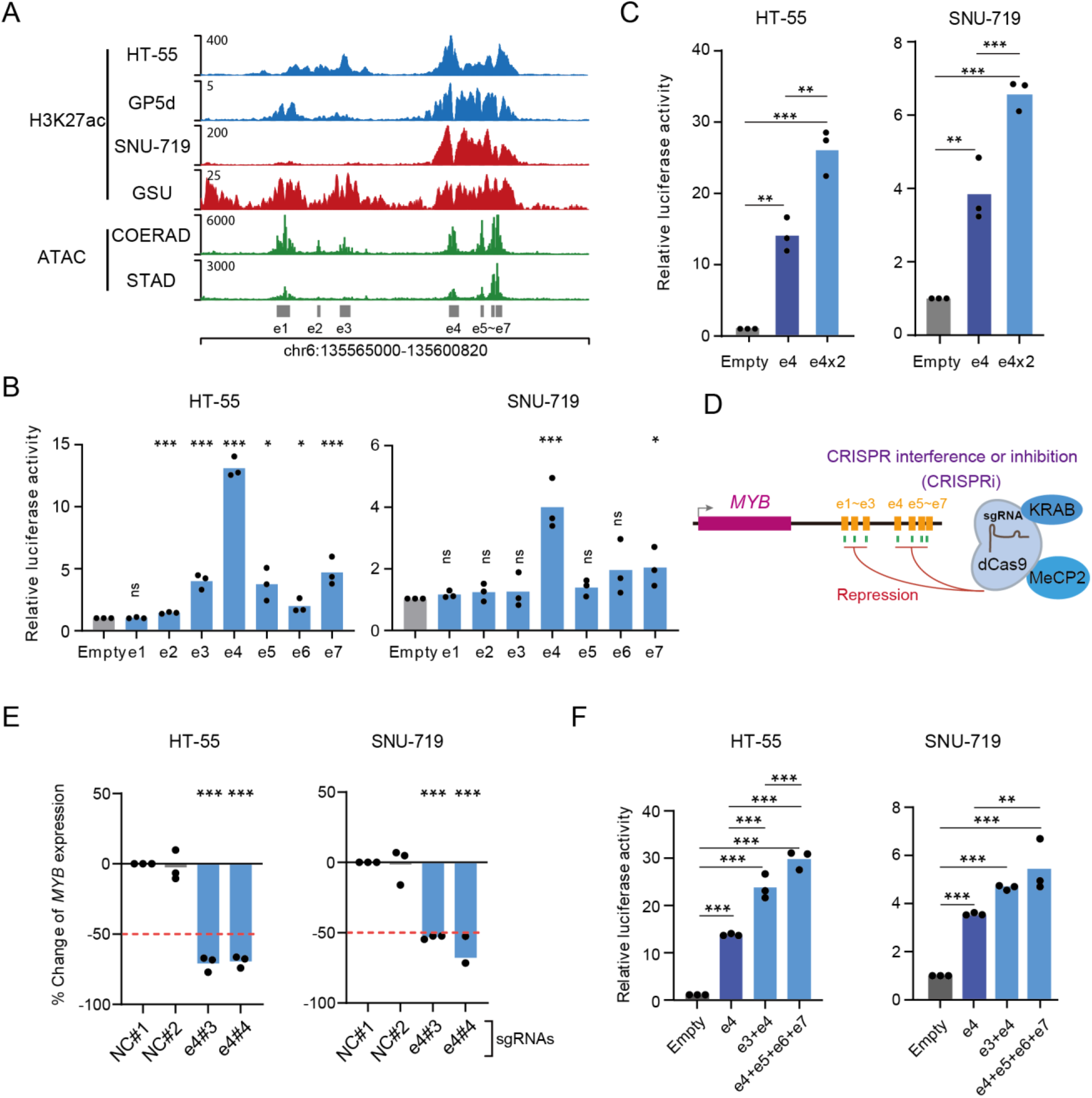
The identification of the predominant enhancer driving MYB expression. **A.** H3K27ac ChIP-seq signal from gastrointestinal adenocarcinoma cancer cell lines and ATAC-seq signal from tumor samples revealed the constituent enhancers e1-e7 for the *MYB*-SE; **B.** Luciferase reporter assay measuring the activity of candidate enhancers in HT-55 and SNU-719 cells. The empty pGL3-promoter vector was used as negative control. N = 3; **C.** Luciferase reporter assay measuring the activity of single or duplicated enhancer e4 (2×e4) in HT-55 and SNU-719 cells. The empty pGL3-promoter vector was used as negative control. N = 3. The P value was determined by One-way ANOVA; **D.** Schematic diagram of the strategy to validate the predominant enhancer by CRISPRi; **E.** The expression of *MYB* upon CRISPRi - mediated repression of enhancer e4. Two separate sgRNAs, as sg-e4#3 and sg-e4#4, were used to target the enhancer e4. NC, non- targeted control. N = 3; **F.** Luciferase reporter assay measuring the activity of single or combinational enhancer in HT-55 and SNU-719 cells. The empty pGL3-promoter vector was used as negative control. N = 3. The P value was determined by One-way ANOVA. *: P ≤ 0.05; **: P ≤ 0.01; ***: P ≤ 0.001.

In advances, we tested the function of e4 for the expression of *MYB* by manipulating its transcriptional activities endogenously. We performed the an improved CRISPR-based interference (CRISPRi, see the details in the Methods and Supplementary Methods)^17, 51^ to suppress the proposed constituent enhancers in four *MYB*-high gastrointestinal adenocarcinoma cell lines (**Fig. 3D**). The results revealed that the repression of e4 resulted in significant reduction in *MYB* expression in all four cell lines, while the suppression of the other six enhancers only had minimal effects (**Supplementary Fig. 4F**). Moreover, these effects of CRISPRi targeting e4 on *MYB* expression were attenuated in the *MYB*-high/-independent CCK-81 cells (**Supplementary Fig. 4F**). The importance of e4 for *MYB* expression was validated by the application of the other two sgRNAs in HT-55 and SNU-719 cells (**Fig. 3E**). In contrast, the activation of e4 by CRISPR-based activation assay (CRISPRa, see the details in the Methods and Supplementary Methods)^52^ in AGS and HGC-27 cell lines, two *MYB*-low/-independent cell lines, dramatically increased the expression of *MYB* (**Supplementary Fig.4G**). These results suggested that e4 of the *MYB*-SE is the predominant enhancer driving *MYB* expression.

It is widely accepted that one enhancer may control the expression of multiple genes within a similar genomic region. Based on the analysis of H3K27ac HiChIP data, we found that the HiChIP anchor of e4-7 has the strongest physical interactions with the *MYB* promoter in all five *MYB*-high and -dependent cell lines (**Supplementary Fig. 5A**). However, we also found four additional genes, *HBS1L*, *BCLAF1*, *MAP7* and *IL20RA,* whose promoters interacted with e4-7 in more than one cell line (**Supplementary Fig. 5A**). We therefore investigated whether e4 controls the transcription of genes other than *MYB*. On one hand, the repression of e4 by CRISPRi had minimal effects on the expression of *HBS1L*, *BCLAF1*, *MAP7*, and *IL20RA* as well as other genes surrounding *MYB* in two cell lines (**Supplementary Fig. 5B-C**). Although the expression of *AHI1* surrounding *MYB* was modestly affected, no interaction with e4 was identified by the analysis of H3K27ac HiChIP data (**Supplementary Fig. 5D-E**).

The enhancers within a SE may work together to regulate the expression of one gene. Intestinally, we also observed the interactions among the multiple enhancers of *MYB*-SE (**Supplementary Fig. 5F-G**). The combination of e4 with other constituent enhancers of *MYB*-SE exhibited higher activity than e4 alone by luciferase assay (**Fig. 3F**), indicating a synergistic effect of the enhancers within *MYB*-SE on transcriptional regulation. Collectively, our findings revealed the predominant role of e4 in the *MYB*-SE in controlling the expression of *MYB*.

### HNF4A controls the activity of e4 for the *MYB*-SE via direct interactions

After confirming the predominant role of e4 for *MYB*-SE, we then explored the potential transcription factors (TFs) occupying the e4 region and controlling its activity. Based on the analysis with phastCons scores^53^ and DNase I hypersensitive^54^, we found that the e4 region exhibited high DNA sequence conservation and a chromatin-accessible state across numerous species (**Fig. 4A**), making it feasible for the downstream analysis. The motif analysis was performed based on publicly available ATAC-seq datasets of tumor samples from 81 patients with primary COREAD and 41 patients with primary STAD from TCGA^48^. Multiple potential transcription factor families which may bind to e4 were identified (**Fig. 4A and Supplementary Fig. 6A**) by the employment of software FIMO^55^ and TRAP^56^ (see the details in the Methods and Supplementary Methods).

**Figure 4.**
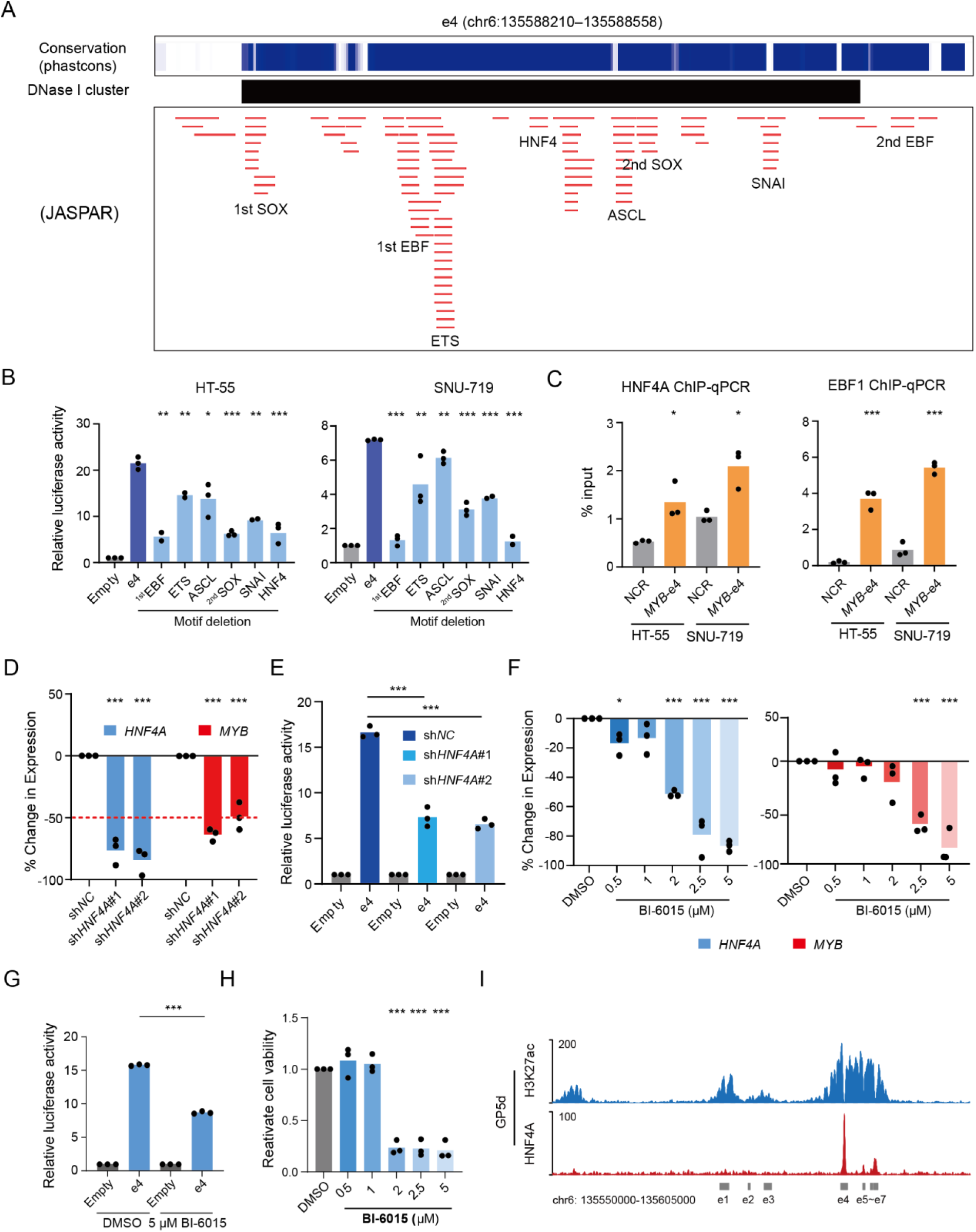
Discovery of the transcription factors controlling the expression of MYB via direct interactions with the predominant enhancer. **A.** The motif analysis of the enhancer e4 sequence. Top: phastCons scores (0:1 range) representing how conserve the enhancer e4 DNA sequence was. Middle: The cluster of DNase I hypersensitive sites demonstrating the accessibility of the enhancer e4. Bottom: The results of JASPAR based prediction for transcription factors’ binding motifs in the enhancer e4; **B.** Luciferase reporter assay measuring the activity of the enhancer e4 upon the deletion of the binding motifs. N = 2-3; **C.** ChIP-qPCR analysis indicating the enrichment of transcription factors HNF4A and EBF1 at the enhancer e4. NCR: Negative control region. N = 3; **D.** The expression of *HNF4A* and *MYB* upon HNF4A knockdown by shRNAs. NC, non-targeted control. N = 3; **E.** Luciferase reporter assay measuring the activity of the enhancer e4 upon HNF4A knockdown in HT-55 cells. N = 3; **F.** The expression of *HNF4A* and *MYB* upon the treatment of HNF4A antagonist BI-6015 in HT-55 cells. N = 3; **G.** Luciferase reporter assay measuring the activity of the enhancer e4 following the treatment of HNF4A antagonist BI-6015 in HT-55 cells. N = 3; **H.** Cell viability of HT-55 cells after 48 hours treatment of BI-6015, measured by Cell Counting Kit-8 (CCK-8); **I.** ChIP-seq tracks for H3K27ac and HNF4A in the *MYB*-SE associated loci in GP5d cells. *: P ≤ 0.05; **: P ≤ 0.01; ***: P ≤ 0.001.

To test whether the expression of *MYB* can be regulated by the binding of a particular TF, we performed genome editing with CRISPR-Cas9 system to disrupt DNA motifs within e4. We found that disruption of the motifs associated with the HNF4, EBF, ASCL, SOX or SNAI families led to remarkable reduction in *MYB* expression (**Supplementary Fig. 6B-C**). Deletion of specific TF-binding motif sequences within e4 revealed that the ASCL, EHF, EBF, HNF4, SNAI, and SOX motifs are important for maintaining e4 activity in the luciferase assay (**Fig. 4B and Supplementary Fig. 6D**). Especially, the depletion of EBF or HNF4 motif had the most dramatic effects on e4 activity.

Based on the results of motif analysis and genome editing, we then focused on two TFs, EBF1 (EBF family motif) and HNF4A (HNF4 family motif), for subsequent analysis. First, binding of EBF1 or HNF4A to e4 was validated by ChIP-qPCR assay in HT-55 and SNU-719 cells (**Fig. 4C**). Importantly, knockdown of *HNF4A*, but not *HNF4G*, reduced the expression of *MYB* and potently inhibited the activity of e4 in the luciferase assay (**Fig. 4D-E and Supplementary Fig.6E-F**). In contrast, knockdown of *EBF1* did not change the expression of *MYB* (Supplementary Fig.7G). Pharmacological inhibition of HNF4A with its antagonist with BI-6015 also suppressed the expression of *MYB* and e4 activity in the luciferase assay (**Fig. 4F-G**). Interestingly, the treatment of BI-6015 also prevented the growth of HT-55 cells (**Fig. 4H**). We further validated that HNF4A binds to e4, e5, e6 and e7 in GP5d cells based on the analysis of published data for HNF4A ChIP-seq^42^. Among these, the binding peak at e4 showed the highest signal (**Fig. 4I**). In brief, these results suggested that HNF4A can bind to e4 and regulate *MYB* expression by regulating the activity of e4 of *MYB*-SE.

### MYB activates cancer-related genes in gastrointestinal adenocarcinomas

To systematically investigate the regulatory functions of MYB in gastrointestinal adenocarcinoma, we preformed ChIP-seq analysis for MYB in HT-55 (a COREAD cell line) and SNU-719 (a STAD cell line) cells. The canonical binding motif of MYB was identified by the software HOMER^57^ and presented at **Fig. 5A**. We also identified the motifs for multiple TFs in MYB-binding DNA sequences, including AP1, KLF5, ETS1, FOXA1 and other cancer-related transcription factors (**Supplementary Fig.7A**), suggesting that these TFs may work together with MYB for transcriptional regulation. We observed that MYB mainly bound to the promoter and distal intergenic regions (**Fig. 5B-C**), suggesting that MYB had the potential to bind regulatory elements including enhancers and SEs.

**Figure 5.**
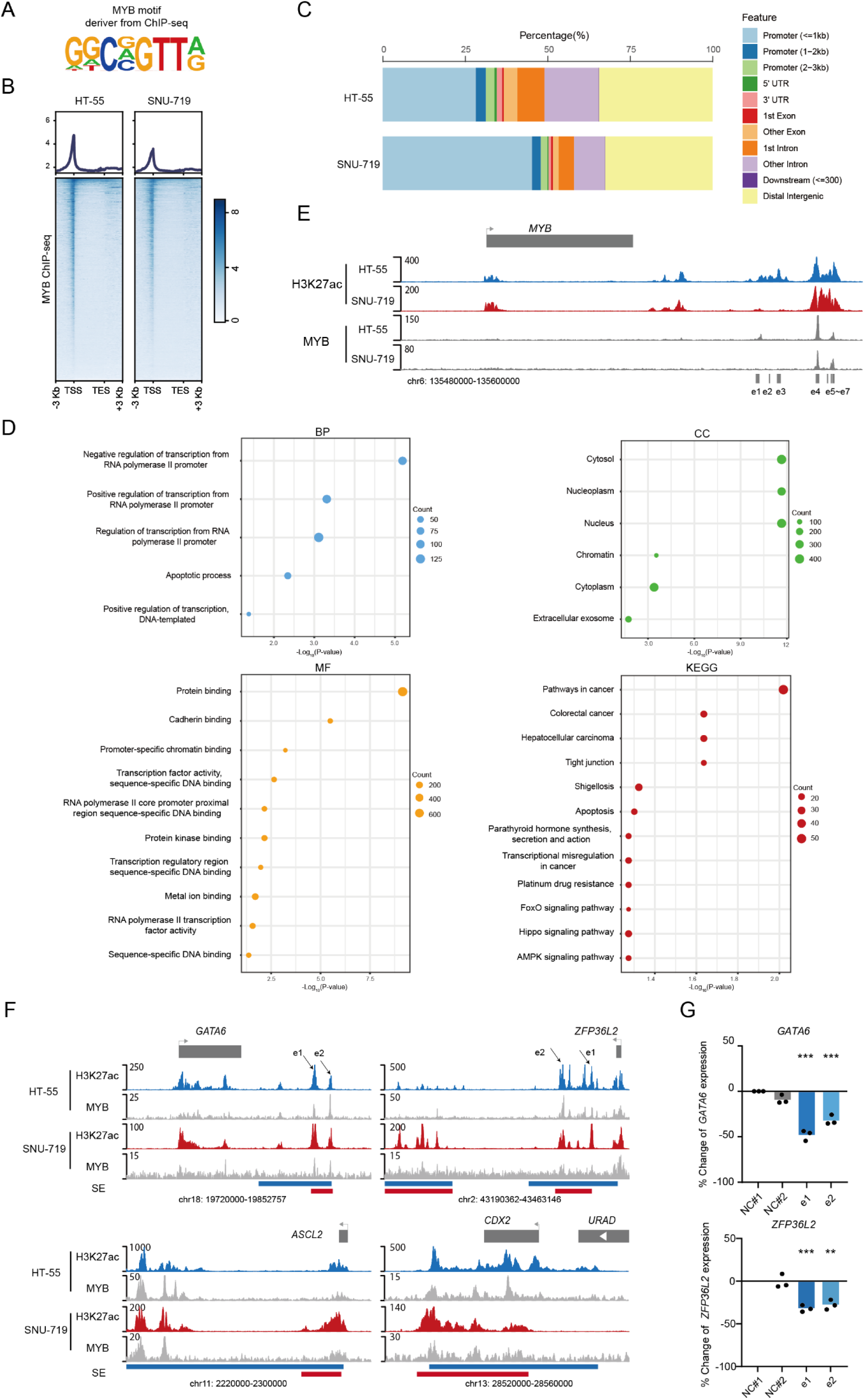
MYB contributes to the high expression of cancer - related genes in gastrointestinal adenocarcinoma. **A.** The prediction of MYB’s DNA binding motifs based on the MYB ChIP-seq data; **B.** The heatmaps showing the distribution of MYB binding sites cross the whole genome in HT-55 and SNU-719 cells. Genes in rows were sorted in decreasing order by the signal intensity. TSS, transcription start site. TES, transcription end site; **C.** Bar charts illustrating the types of MYB binding sites in HT-55 and SNU-719 cells; **D.** Functional enrichment analysis of the nearest genes to the MYB binding sites; **E.** ChIP-seq tracks of MYB in HT-55 and SNU-719 cells showing that MYB bound to its super-enhancer; **F.** MYB bound to the candidate super-enhancers of various cancer-related genes; **G.** Validating the functions of the candidate super-enhancers for *GATA6* and *ZFP36L2* by CRISPRi. N = 3. *: P ≤ 0.05; **: P ≤ 0.01; ***: P ≤ 0.001.

Assigning the MYB binding sites to their nearest genes discovered a total of 1240 genes in both HT-55 and SNU-719 cells (**Supplementary Fig. 7B and Supplementary Table 3**). The enrichment of GO and KEGG terms revealed that these genes were involved in apoptosis, cancer development and transcription control (**Fig. 5D**). Specifically, we found that MYB bound to e4, e6 and e7 of *MYB*-SE based on the analysis of ChIP-seq and ChIP-qPCR (**Fig. 5E and Supplementary Fig. 7C**). Knocking down *MYB* significantly inhibited the activity of e4 in the luciferase assay (**Supplementary Fig. 7D**), indicating that MYB can regulate the transcriptional activity of e4. Thus, an auto-regulatory loop, which is a typical feature of lineage-specific oncogenes^58–63^, may exist between MYB and *MYB*-SE. In addition, we also found that MYB can bind to the SEs of other cancer-related genes (**Fig. 5F and Supplementary Fig. 7E**). Specifically, CRISPRi-based suppression of MYB binding sites in HT-55 cells downregulated the expression of *GATA6* and *ZFP36L2*, two well-known genes with oncogenic roles in gastrointestinal adenocarcinoma^16, 64^ (**Fig. 5G**).

To investigate the genes whose expression was regulated by MYB, we performed RNA sequencing (RNA-Seq) analysis for the *MYB*-depleted HT-55 cells. Differential expression analysis identified 532 upregulated genes and 496 downregulated genes in the *MYB*-depleted cells comparing to cells with scrambled control (**Supplementary Fig. 8A**). Integrative analysis of RNA-seq and ChIP-seq data revealed 407 genes which were the potential direct targets of MYB (**Supplementary Fig. 8B and Supplementary Table 4**). Gene set enrichment analysis (GSEA) showed that MYB targeted genes, which were downregulated by *MYB*-depletion, were related to Notch signaling (**Supplementary Fig. 8C**). These findings indicated that MYB directly controlled the expression of multiple cancer related genes, including itself, which may contribute the development of gastrointestinal adenocarcinoma.

### Targeting the e4 of *MYB*-SE suppresses the growth of gastrointestinal cancer cells in vitro and in vivo

Considering the significance of MYB for the transcription of *MYB* and multiple oncogenes, we hypothesized that *MYB* is required for the high proliferation rate of gastrointestinal cancer cells. To validate this, we first used an shRNA system to knock down *MYB* in HT-55 and SNU-719 cells. We observed that suppressing the expression of *MYB* significantly inhibited the cell growth (**Fig. 6A and Supplementary Figure 9A-B**). To characterize the effects of the e4 on cellular function, we suppressed its activity by CRISPRi system and discovered a significant decrease of cell growth as well (**Fig. 6B, Supplementary Fig. 9C**). Additionally, the suppression of e4 prevented the formation of colonies (**Fig. 6C**), and reduced the positive rate of EdU staining as the marker for cell proliferation in HT-55 cells (**Fig. 6D**). Mechanistically, the suppression of *MYB* downregulated the expression of genes related to Notch signaling (**Supplementary Fig. 9D**). Particularly, the treatment of Notch antagonist BMS-986115 prevented the growth of HT-55 cells (**Fig. 6E**).

**Figure 6.**
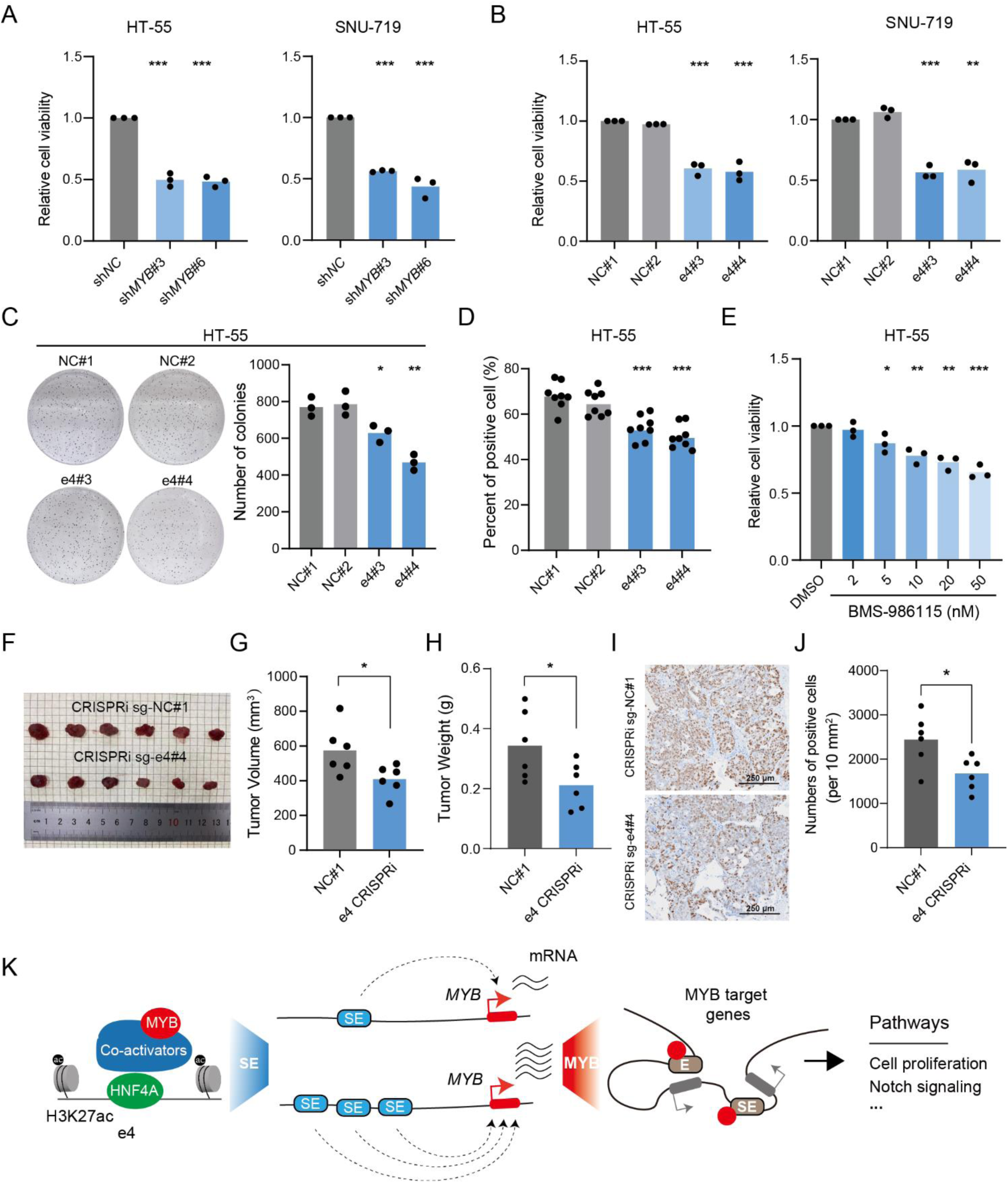
The predominant enhancer of MYB as a master regulator for the development of gastrointestinal adenocarcinoma. **A-B.** Cell viability of HT-55 and SNU-719 cells upon *MYB* knockdown **(A)** or repression of the enhancer e4 **(B)**, measured by Cell Counting Kit-8 (CCK-8). N = 3; **C.** The effects of repression of the enhancer e4 on colony formation of HT-55 cells. N = 3; **D.** The effects of repression of the enhancer e4 on cell proliferation of HT-55 cells as measured by EdU staining. N = 8; **E.** Cell viability of HT-55 cells upon the treatment of Notch antagonist BMS-986115, measured by Cell Counting Kit-8 (CCK-8). N = 3; **F-H.** The growth of the xenografts based on HT-55 cells were significantly inhibited after repression of the enhancer e4. N = 6; **I.** The representative pictures of Ki-67 staining of xenografts. Scale bar, 250 µm; **J:** The average number of Ki-67 positive cells per 10 mm^2^ xenograft tumor tissue. N = 6; **K.** Schematic diagram: The expression of *MYB* gene was activated by super-enhancer amplifications. The overexpressed MYB protein directly upregulated its target genes and promoted the development of gastrointestinal adenocarcinoma. **: P ≤ 0.05; **: P ≤ 0.01; ***: P ≤ 0.001.

We then tested the effects of *MYB in vivo* in the xenografts with HT-55 cells of nude mice. We observed that both the volume and weight of tumor xenografts were significantly downregulated by suppressing e4 activity or knockdown the endogenous *MYB* (**Fig. 6F-H and Supplementary Fig. 9E-G**). The expression of Ki-67, a marker of cell proliferation, was decreased in upon e4 inhibition or *MYB* knockdown (**Fig. 6I-J and Supplementary Fig. 9H**). Taken collectively, these data indicated that *MYB* behaved as a potential oncogene to promote the development of gastrointestinal adenocarcinomas.

## Discussion

To decipher the biological and pathological basis of gastrointestinal adenocarcinoma may improve its diagnosis and therapy. By systematically investigating the active enhancer profiles of tumor samples as well as cell lines, we identified a lineage-specific super-enhancer for *MYB* containing a dominant enhancer e4, which was duplicated in gastrointestinal adenocarcinoma. With an auto-regulatory loop, MYB regulates the development of gastrointestinal adenocarcinoma by directly controlling the transcription of multiple carcinogenic genes associated with Notch signaling. These results suggested the specific role of *MYB* in the carcinogenesis of gastrointestinal adenocarcinoma (**Fig. 6K**).

Various types of genomic alterations for *MYB* locus have been detected within its coding region of human cancer, such as duplication, translocation, gene-fusion, and intragenic mutations^21–24^. However, these genomic alterations can only explain the overexpression of *MYB* in a certain portion of cancer. It has been reported that the abnormal activation of enhancer may be a major epigenetic aberration for various types of cancer^14–17, 65–67^. The current study provided a sophisticated analysis for a genetically duplicated SE, which was looped to the *MYB* promoter and controlled its expression specifically in the gastrointestinal adenocarcinoma. Especially, the predominant enhancer e4 interacts with other enhancers of this SE to regulate its transcriptional activities. These results confirmed the existence of “facilitators”, as the elements contributing to the classical enhancers’ function without inherent activities^68^. It is possible that some of low-activity enhancers are the facilitators for the e4 of *MYB*-SE.

The MYB protein is a key therapeutic target for multiple types of cancer, such as T cell acute lymphoblastic leukemia (T-ALL), breast cancer, prostate cancer, et al^19^. The current study expanded our knowledge about MYB by elucidating its important role in the development of gastrointestinal cancer. We found that MYB bound to the e4 of the *MYB*-SE to regulate the expression of *MYB*, representing a self-regulatory circuit as a feature for lineage-specific master transcription factors^58–63^. We also found that HNF4A controlled the transcription of *MYB* as well as the growth of gastrointestinal adenocarcinoma via binding to the e4 of the *MYB*-SE. As a mRNA-based strategy has been developed to target HNF4A in preclinical models for liver fibrosis^69^, similar strategy can be used to control the development of gastrointestinal adenocarcinoma with HNF4A dependent overexpression of *MYB*.

In addition to its functions in regulating the progress of cancer, *MYB* also plays a role in the developmental process of neural tissues, colonic crypts, breast tissue as well as hematopoietic system^70–74^. We show that the suppression of e4 in *MYB*-SE, which downregulated the expression of *MYB*, was able to decrease the growth of gastrointestinal adenocarcinoma *in vitro* and *in vivo*. This finding highlighted the significance of e4 for the function of *MYB*, which may inspire further studies about the *MYB* associated non-coding variants in the development of non-malignant tissues in a population level. Particularly, the single nucleotide polymorphism (SNP) or structural variation (SV) of e4, identified by genome-wide association study (GWAS) in ancient or modern human, needs to be revisited to elucidate its role in human evolution.

In summary, our study revealed the existence of a *MYB* associated lineage-specific super-enhancer, containing a predominant enhancer e4 and a few potential facilitators, which controlled the development of gastrointestinal adenocarcinoma. These findings underscore the potential of non-coding elements, such as super-enhancers, as viable therapeutic targets for cancer treatment.

## Materials and Methods

### Cell lines and cell culture

AGS and HGC-27 cells were obtained from the Cell Bank of Chinese Academy of Sciences. HT-55, GP2d and CCK-81 cells were obtained from Nanjing Cobioer Biosciences company. HEK293-FT and SNU-719 cells were gifted by Lin Deng, Shenzhen Bay Laboratory. AGS, HGC-27 and SNU-719 cells were cultured in RPMI 1640 medium (Gibco, #11875119) supplemented with 10% FBS (Gibco, #10099141C) and 1% penicillin-streptomycin (Gibco, #10378016). HEK293-FT, HT-55, GP2d and CCK-81 cells were cultured in DMEM high glucose medium (Gibco, #11995073) supplemented with 10% FBS and 1% penicillin-streptomycin. All cells were verified by short tandem repeat analysis and tested negative for mycoplasma using the Mycoplasma qPCR Detection Kit (Beyotime, #C0303S).

### Luciferase reporter assay

Luciferase reporter assays were performed as previously described^14^. Firstly, enhancer e4 was amplified from the HT-55 cell’s genomic DNA, then the enhancer e4 regions were cloned upstream of the pGL3-promoter vector (Promega, #E1761) using XhoI (NEB, #R0146S) and MluI (NEB, #R3198S) restriction enzyme sites. For cloning the 2×e4, e4 region was cloned using XhoI and MluI restriction enzyme sites, followed the another e4 region was cloned using KpnI (NEB, #R3142S) and MluI restriction enzyme sites. For the combination of e4 and other constituent enhancers, other enhancers were cloned using XhoI and MluI restriction enzyme sites into 2×e4 vector. The enhancer luciferase reporter constructs were co-transfected with a control luciferase construct, named *Renilla* (Promega, #E2271), into cells using PEI (Polysciences, #24765) or lipofectamine 3000 (Invitrogen, #L3000015). The luciferase signal was first normalized to the *Renilla* luciferase signal and then normalized to the signal from the empty pGL3-promoter vector. Primers used for cloning are listed in **Supplementary Table 5**.

### CRISPR-mediated DNA motif cutting, enhancer repression and activation

According to the ATAC-seq peaks within the candidate *MYB*-SE, the DNA sequence of the enhancers was obtained. Then, CRISPR/Cas9 sgRNAs were designed using the CRISPick tool from the Broad Institute^52, 75^ and negative control, non-targeting sgRNAs were used as previously reported ^17^. For DNA motif cutting, cells were first infected with lenti-Cas9-blast vector (Addgene, #52962) and selected with 10ug/ml of blasticidin (Beyotime, #ST018-5ml) for at least 7 days. Cells stably expressing Cas9 then subsequently infected with LentiGuide-Pour vector (Addgene, #52963) carrying either non-targeting sgRNAs or sgRNAs targeting the DNA motif and selected with 2ug/ml puromycin (Beyotime, #ST551-250mg) for at least 5 days before any molecular or cellular assays. For enhancer repression, cells were first infected with lenti-dCas9-KRAB-MeCP2-blast vector (Addgene, #122205) and selected with 10ug/ml of blasticidin for at least 7 days. Cells stably expressing dCas9-KRAB-MeCP2 then subsequently infected with LentiGuide-Pour vector carrying either non-targeting sgRNAs or sgRNAs targeting the *MYB* enhancers and selected with 2ug/ml puromycin for at least 5 days before any molecular or cellular assays. For enhancer activation, cells were first infected with lenti-dCas9-VP64-blast vector (Addgene, #61425) and selected with 10ug/ml of blasticidin for at least 7 days. Cells stably expressing dCas9-VP64 then subsequently infected with pXPR-502 vector (Addgene, #96923) carrying either negative control sgRNAs or sgRNAs targeting the enhancer e4 and selected with 2ug/ml puromycin for at least 5 days before any molecular or cellular assays. All sgRNA sequences are listed in **Supplementary Table 5**.

### Cell growth assays and colony formation assays

For cell growth assays, selected cells were collected. 5000 cells were seeded into 96-well plates (100 µL/well). After 5 days, 10 µL of Cell Counting Kit-8 (CCK-8) (Beyotime, #C0040) was added into each well according to the manufacturer’s instructions and incubated for 1 h. Optical density was measured at 450 nm using a microplate reader. For colony formation assays, 5000 cells were seeded into 6-well plates (2 mL/well). The cell culture medium was changed every three days. After culture for 18 days, cells were fixed with methanol (100%) and stained with crystal violet solution (25% methanol and 75% crystal violet) for 20 min at room temperature.

### *In vivo* xenograft tumor assays

All animal experiments were performed in accordance with procedures approved by the Fudan University ethical committee. All the mice were maintained under a 12 hr light/12 hr dark cycle at constant temperature (23 °C) and specific pathogen-free conditions with free access to food and water. A total 1 million HT-55 cells with or without CRISPRi system mediated e4 repression or shRNA system mediated *MYB* knock-down were resuspended in 100 μL serum-free DMEM and supplement with 100 μL Matrigel (Coring, #FAL-354248) (total volume 200 μL). Then the cells were subcutaneously injected into the flanks of female nude mice (BALB/c, Nu/Nu, 5-6 weeks old, female from GemPharmatech). Tumor growth was examined every 4–5 days, and tumor length and width were measured using calipers. Until tumor volume reached approximately 1000 ∼ 1500 mm^3^(the tumor size not exceeded 2 cm in any dimension), xenograft tumor-bearing mice were sacrificed, the tumor xenografts were isolated and weighted. Tumor volume was calculated suing the following formula: (length × width^2^) × 0.52.

### Public data usage

This study used TCGA publicly available ATAC-seq data were downloaded from NCI Genomic Data Commons data portal (UTR: https://gdc.cancer.gov/about-data/publications/ATACseq-AWG), publicly PCAWG SV data were downloaded from PCAWG data portal(UTR: https://dcc.icgc.org/releases/PCAWG/consensus_sv), The remaining publicly available data were downloaded from GEO. Accession numbers of public dataset used in this study listed in Supplementary Table 2.

### Quantification and statistical analysis

All statistical comparisons between two groups were performed by GraphPad Prism software 9.0 using a two-tailed unpaired *t*-test, unless specified. The variance between the statistically compared groups was similar.

## Data availability

The MYB ChIP-seq and RNA-seq datasets in this study have been deposited in the Gene Expression Omnibus (GEO) database with accession number GSE234204. The rest of the data supporting this study are available in the article or Supplementary Information.

## Author contributions

Conceptualization, F.L., S.W., L.C., N.J., X.C. and J.L.; Investigation, F.L., S.W., L.C., N.J., X.C. and J.L.; Analysis, F.L., S.W., L.C., N.J., X.C. and J.L.; Writing, F.L., S.W., L.C., N.J., X.C. and J.L.; Data Visualization, F.L., S.W., L.C., N.J., X.C. and J.L.; Funding Acquisition, X.C. and J.L.; Supervision, X.C. and J.L..

