## Supplementary figures for "Systemic genome-epigenome analysis captures the lineage specificity and functional significance for *MYB* associated super-enhancer in gastrointestinal adenocarcinoma"

#### Supplementary Materials and Methods

##### Analysis of PCAWG and TCGA datasets

The list of gastrointestinal adenocarcinoma structural variants was downloaded from the Pan-Cancer Atlas of Whole Genomes (PCAWG) project (URL:https://dcc.icgc.org/releases/PCAWG/consensus_sv)[1]. Continuous 5 kb bins were tiled across the entire genome using BEDTools[2], and their duplication frequency in gastrointestinal adenocarcinoma were calculated. The adjacent bins were stitched together to form duplications and amplicons (duplication frequency ≥ 5). TCGA ATAC-seq data was download from the NCI Genomic Data Commons data portal (URL: https://gdc.cancer.gov/about-data/publications/ATACseq-AWG)[4]. We used the normalized bigWig data for Integrative Genomics Viewer presentation and we converted the DNA coordinates of the ATAC sites from hg38 to hg19 using the liftOver tool[5]. For RNA expression analysis, we used GEPIA2[6] for analyzing the RNA sequencing expression data of tumors and normal samples from the TCGA and the GTEx projects.

##### DepMap CRISPR screen analsyis

We analyzed the publicly available CRISPR screen results on cancer cell lines from the Broad Institute Dependency Map (DepMap) project[7]. The Spearman correlation between RNA expression results from the Cancer Cell Line Encyclopedia (CCLE) project and CERES, the CRISPR dependency score, was calculated as previously reported[8]. Also, we compared the correlation between the Broad CRISPR screen results and the Sanger Institute CRISPR screen results or the Broad shRNA screen results. Gene-level copy number data that is $\log_2$ transformed with an absolute-count of 1; $\log_2$(copy number/2). Inferred from WGS, WES or SNP array depending on the availability of the data type. All data were downloaded in September of 2023.

##### Motif enrichment analysis

ATAC-seq signal from COREAD and STAD was to narrow down DNA coordinates of enhancer e4 region. With the default parameters, the FIMO software[9] was used to identify transcription factor binding motifs from the JASPAR motif database[10] that are present in the enhancer e4. Also, accessible chromatin region of e4 was subject to motif analysis using Transcription Factor Affinity Prediction (TRAP) web tools[11]. The region was analyzed using JASPAR vertebrates as the matrix file, all human promoters as the background model, and Benjamini-Hochberg as the multiple test correction. We generated ChIP-seq data for MYB in HT-55 and SNU-719 cells in the present work. HOMER[12] was used to analysis the motifs, HOMER script "findMotifsGenome.pl" was then performed for motif search with parameters –no motif and hg19. Besides, deeptools[13] (v3.5.1) and R package ChIPseeker[14, 15] (v1.32.1) were used to annotate the ChIP-seq results.

**Identification of predicted super-enhancers**

H3K27ac ChIP-seq datasets were downloaded from the Gene Expression Omnibus (GEO) dataset. H3K27ac ChIP-seq fastq data were aligned to the hg19 genome assembly using Bowtie2[16] (v2.2.5). Peaks were called using MACS2[17] (v2.2.7.1). BedGraph files were generated by MACS2, sorted, and normalized by the mean genome-wide bedGraph signal in each dataset. Bigwig tracks were generated from bedGraph using UCSC tool bedGraphToBigWig, and visualized by gTrack (v0.1.0) (https://github.com/mskilab-org/gTrack). Predicted enhancers were identified as enriched H3K27ac regions at least 2.5kb away from annotated transcription starting site (TSS) regions. We allowed enhancers within 15,000 bp to be stitched together. Predicted enhancers were then further divided into super-enhancers and typical enhancers using the ROSE[18] algorithm.

**Analysis of HiChIP and Hi-C data**

HiChIP and Hi-C data was downloaded from the Gene Expression Omnibus (GEO) database. The sequencing reads were aligned to the hg19 human genome using the HiC-Pro pipeline[19], we the used the AllValidPairs output from HiC-Pro that lists all valid paired-end tags (PETs). Then, we used the hichipper pipeline[20] to call significant chromatin loops and used TCGA ATAC sites identified in COREAD and STAD cancer as pre-determined peaks for loop calling by hichipper. The loops supported by at least two PETs and associated with an FDR value $< 0.05$ were considered as significant loops. We next mapped the PETs to continuous 5kb bins within the regions of interest and presented the interactions as contact map using the gTrack R package.

**Super-enhancers nominated genes analysis**

The expression level of Super-enhancers nominated 205 protein-coding genes analysis by GEPIA2[6] tools, including TCGA and GTEx project data. Gene Ontology (GO), KEGG and Reactome analysis were performed using the online DAVID tool[21]. Using H3K27ac HiChIP data in HT-55 and SNU-719 cell lines for identified the interactions between the super-enhancers and nominated protein-coding genes. Identification of oncogenes based on annotated information from COSMIC (https://cancer.sanger.ac.uk/census), OncoKB (https://www.oncokb.org/cancerGenes) and ONGene (https://ongene.bioinfo-minzhao.org/) databases.

**RNA-seq**

Total cells RNA was extracted using the Zymo Quick-RNA miniprep kit (Zymo, #R1055) according to the manual. PolyA mRNA was purified using NEBNext PolyA mRNA Isolation Module (NEB, #E3370S), as per the manufacturer's instructions. RNA-seq libraries were prepared using the NEBNext Ultra Directional II RNA library prep kit and sequenced by Illumina/BGI with a 150 bp paired-end run configuration. Three biological replicates per sample.

**RNA-seq analysis**

Quality control was performed using the FastQC (v0.11.7) tool and the results were analyzed. Cleaned RNA-seq reads were mapped using STAR[22] (v. 2.7.10) against the human genome (hg19). Uniquely mapped reads were quantified with HTSeq[23] (v.0.11.3) and protein-coding genes with nonzero read count ($n = 20138$) were included for downstream analysis. Differential expression of protein-coding genes was analyzed with these counts using Bioconductor package DESeq2[24] (v.1.40.1). Significantly differential expression was considered by setting $adjP < 0.05$ and $FC \geq 1.5$ between sh*MYB* and shNC. Gene Ontology (GO) and KEGG analysis were performed using the online DAVID tool[21]. The gene set enrichment analysis was performed according to the instructions (https://www.gsea-msigdb.org/gsea/index.jsp).

**ChIP-seq and ChIP-qPCR**

Chromatin-immunoprecipitation followed by massive parallel sequencing (ChIP-seq) was performed as previously described[25]. In brief, 6 million cells were crosslinked with 1% formaldehyde and lysed. The chromatin extract was sonicated by the QSonica Q800R3 sonicator (25 minutes, 70% amplitude) and immunoprecipitated with anti-MYB antibody (Abcam, #ab45150) that were mixed with Dynabeads A and G (Invitrogen, #10002D/10004D). The immunoprecipitated DNAs were then used to construct the ChIP-seq library, following the protocol provided by the manufacturer and sequenced on Illumina Xten with the PE 150 method, and sequenced by Illumina NovaSeq. For chromatin immunoprecipitation (ChIP)-qPCR assays, we designed primers targeting MYB, EBF1 and HNF4A binding sites from e4. We also used sonicated genomic DNA to normalize variance in primer efficiency. All the primers are listed in **Supplementary Table 5**.

**ChIP-seq analysis**

Sequencing reads aligned to the hg19 human genome reference by Bowtie2 (v 2.2.5), and SAMtools[26] (v1.6) was used to sort and index the aligned reads. PCR duplicates were marked and removed using Picard (v2.18.29) MarkDuplicates function. MYB binding sites were called by MACS2 (v2.2.6) using the default parameters (q value $< 0.05$). Bedgraph files generated with MACS2 were later converted to bigwig using UCSC bedGraphToBigWig tool, which was then visualized by Integrative Genomics Viewer (IGV)[27]. The H3K27ac HiChIP data of HT-55 and primary colonic epithelial cells were analyzed by chip-seq pipeline to obtain ChIP-seq-like H3K27ac signal. Gene Ontology (GO) and KEGG analysis of MYB ChIP-seq were performed using the online DAVID[21] tool.

**Genomic STARR-seq analysis**

GP5d STARR-seq data was downloaded from GEO dataset, the accession number is GSM5454434. The analysis was carried out according to the previously reported methods[28]. In brief, the sequencing reads were aligned to the human genome (hg19) using Bowtie2 (v2.2.5, option –maxins 1000). Mapped read pairs were deduplicated using Picard (version 2.9.0) and paired-end reads that had mapping quality <20 or mapped in a discordant orientation were discarded using SAMtools (v1.6). Then, the active enhancers were identified by calling the peaks from the STARR-seq–enriched RNA fragments against the plasmid input sample using MACS2 (v2.2.6, -f BAMPE).

**Site-directed deletion of motif sequence**

The Q5® Site-Directed Mutagenesis Kit (NEB, #E0554) was used to generate deletions of the predicted motif sequences in the e4 region. Primers designed by using the NEB online design software, NEBaseChanger™, and the sequences are listed in **Supplementary Table 5**.

**Plasmid transfection and lentiviral production**

sgRNAs for CRSISPRi and CRSISPR-Cas9-mediated DNA motif cutting were cloned into the BsmBI (Thermo Fisher Scientific, #FD0454) sites of the lentiGuide-Pour (Addgene, #52963) lentiviral vector. sgRNAs for CRSISPRa were cloned into the BsmBI sites of the pXPR_502 (Addgene, #96923) lentiviral vector. The double-stranded oligonucleotide shRNA was cloned into the AgeI/EcoRI (NEB, #R3552L/R3101L) sites of the pLKO.1 (Addgene, #8453) lentiviral vector. All sgRNA and shRNA target sequences were listed in **Supplementary Table 5**. HEK293-FT cells were co-transfected with lentiviral packaging vectors in 10% FBS, 0.1% P/S DMEM high glucose medium. Plasmids transfection was performed according to the manual of CalPhos Mammalian Transfection Kit (Clontech, #631312). 24 h after transfection, the medium was replaced with fresh medium. 48h after transfection, the supernatant containing virus was collected and filtered through a 0.45 μm filter (Merck, #SLHP033R) to remove cells and debris. Virus then was added to cells or stored at -80 °C.

**BI-6015 treatment**

HNF4A antagonist BI-6015 (MedChemEpress, #HY-108469) was dissolved with DMSO into 5 mM storage solution, and stored at -80 °C. When needed, the storage solution was diluted into the corresponding concentration with working solution. The cells were treated for 48 h before tested with various assays.

**BMS-986115 treatment**

BMS-986115 (Notch inhibitor 1) (MedChemEpress, #HY-12860) was dissolved with DMSO into 5 μM storage solution, and stored at -80 °C. When needed, the storage solution was diluted into the corresponding concentration with working solution. The cells were treated for 5-day before tested with various assays.

**Quantitative RT-PCR**

Total RNA was isolated using TRIzol (TAKARA, #9109). 1μg RNA was reverse transcribed into cDNA using LunaScript® RT SuperMix Kit (NEB, #E3010L). qPCR was performed with Luna® Universal qPCR Master Mix (NEB, #M3003E). The fold changes were normalized to *HPRT1* or *R28S*. The primers used for qPCR were listed in **Supplementary Table 5**.

**Western Blot**

Cells were harvested and washed in cold PBS for 3 times. Cells were lysed with NP40 lysis buffer (1% NP40, 150mM NaCl, 50mM Tris-HCl, pH 8.0) supplemented with protease inhibitors and sonicated with QSonica Q800R (pulse: 30s on/ 30s off, sonication time: 3 mins, amplitude: 50%). c-MYB (ProteinTech, #17800-1-AP, 1:1000), HNFA4 (CST, #3113S) and ACTB (CWBio, #CW0096M, 1:2000) antibodies were used.

**Immunohistochemistry and EdU analysis**

4% polyformaldehyde-fixed (Beyotime, #P0099), paraffin-embedded tumor sections (4-μm-thick) were dewaxed and rehydrated through graded alcohol to water before antigen retrieval, followed by treatment with 3% hydrogen peroxide. The sections were incubated with special antibody against Ki-67 (CST, #9449S) at 4°C diluted at 1: 400, subsequently stained with the DAB (Thermo Fisher Scientific, #34002) on the following day. Then, the sections were counterstained with hematoxylin (Beyotime, #C0107). For EdU staining, 10000 cells were seed into 96-well plate and cultured overnight. Then Staining analysis was performed using Cell-Light EdU Apollo488 In Vitro Kit (RIBOBIO, #C10310-3).

**Supplementary Figure Legends:**

**Supplementary Figure 1. Enrichment and functional analysis of the common SEs nominated protein-coding genes**

**A.** Enrichment and functional analysis of 205 SEs nominated protein-coding genes. MF: molecular function. CC: cellular component; **B.** Reactome analysis for 205 SEs nominated protein-coding genes; **C.** The expression of 44 genes upregulated in tumor samples are presented in heatmap.

**Supplementary Figure 2. The gastrointestinal specificity for *MYB*-SE**

**A.** Venn diagram showing the overlap of oncogenes associated to SEs and upregulated genes with physical interactions to SEs in gastrointestinal adenocarcinoma. **B.** Venn diagram of the shared SEs associated genes between gastrointestinal cancer cell lines and tumor samples; **C.** ChIP-seq signal tracks showing the enrichment of H3K27ac at the *MYB*-SE loci. The blue denoted COREAD tracks and red denoted STAD tracks; **D.** H3K27ac ChIP-seq tracks at the *MYB* loci in cell lines representing each cancer type. ESAD: esophageal adenocarcinoma; BLCA: bladder cancer; PADD: pancreatic adenocarcinoma; LIHC: liver hepatocellular carcinoma; UCEC: uterine corpus endometrial carcinoma; LUAD: lung adenocarcinoma; BRCA: breast cancer; PRAD: prostate adenocarcinoma; LAML: acute myeloid leukemia; **E.** ChIP-seq signal tracks of H3K27ac at the *Myb* and *Myc* loci in mouse CRC cell lines; **F.** The expression of *Myb* and *Myc* level in mouse CRC cell lines, data from RNA-seq; **G.** The expression of *MYB* in cancer tissues and normal tissues from the TCGA and GTEx database. The results were calculated by Gene Expression Profiling Interactive Analysis 2 (GEPIA2); **H.** The expression of *MYB* in the COREAD or STAD samples and pair-matched normal tissues. The value for the y-axis is $log_2$ transformed (count+1); **I.** The correlation calculated by spearman's rank correlation coefficient between *MYB* expression and *MYB* gene copy number based on the data from CCLE. NS: no significance. ***: P $\leqslant$ 0.001.

**Supplementary Figure 3. The chromatin landscape of the MYB locus in gastrointestinal adenocarcinoma**

**A.** The expression level and dependency score of *MYB* in the indicated cell lines based on DepMap; **B.** The integration of H3K27ac ChIP-seq, HiChIP, and Hi-C data showing three-dimensional structure of the *MYB* loci in HCT 116, RKO, AGS and PCE cells. PCE: normal human primary colonic epithelial cells. Turquoise box showed that there was no interaction between *MYB* promoter and *MYB*-SE; **C.** H3K27ac ChIP-seq tracks at the *MYB* loci in *MYB*-high/-dependent, *MYB*-high/-independent and *MYB*-low/-independent gastrointestinal adenocarcinoma cancer cell lines. The copy number status of *MYB* gene is indicated under the name of the cell lines.

**Supplementary Figure 4. The enhancer regulates the expression of *MYB***

A. The correlation of *MYB* dependency scores from DepMap versus the results from Sanger Institute CRISPR screen (left) or the Broad Institute shRNA screen (right); **B.** The correlation between expression and dependency score of *MYB* across all the cancer cell lines in DepMap. The results of COREAD and STAD cell lines were highlighted in red and blue respectively; **C.** The H3K27ac ChIP-seq track, ATAC-seq track and STARR-seq track at the *MYB*-SE locus of GP5d cells. Note that e4 has the strongest STARR-seq signal; **D.** Luciferase reporter assay measuring the activity of candidate enhancers activity in HEK293-FT cells. The empty pGL3-promoter vector was used as negative control. N = 2; **E.** Schematic representation of enhancer e4 duplication in pGL3-promoter vector; **F.** The effects of CRISPRi on the candidate enhancers for *MYB* expression in GP2d, CCK-81, HT-55 and SNU-719 cells. N = 3; **G.** The effects of CRISPRa on the enhancer e4 for *MYB* expression in two *MYB*-low/-independent cell lines AGS and HGC-27. N = 2. *: P ⩽ 0.05; **: P ⩽ 0.01; ***: P ⩽ 0.001.

**Supplementary Figure 5. The enhancer e4 is predominant for *MYB*-SE**

**A.** Number of paired-end tags (PETs) that connecting e4-7 anchor and the candidate genes' promoter in five cell lines based on H3K27ac HiChIP data; **B-C.** The expression of candidate genes with CRISPRi on enhancer e4 in HT-55 **(B)** or SNU-719 **(C)** cells. The genes were annotated with the same color as **A**; **D-E**. The expression of genes adjacent to *MYB* with CRISPRi on enhancer e4 in HT-55 **(D)** or SNU-719 **(E)** cells. The genes were annotated with the same color as **A**; **F.** The contact heatmap based on H3K27ac HiChIP data revealed that enhancer e4 interacted with other candidate enhancers within *MYB*-SE in HT-55, GP5d and SNU-719 cells; **G.** Number of PETs that connecting e4-7 anchor and e1-3 or e1 anchor based on H3K27ac HiChIP data in five cell lines. *: P ⩽ 0.05**; P ⩽ 0.01; ***: P ⩽ 0.001.

**Supplementary Figure 6: Candidate transcription factors regulating MYB expression via interactions with enhancer e4**

**A.** The transcription factors which may bind to enhancer e4 based on the prediction of TRAP motif analysis; **B.** Demonstration of the motifs to be disrupted by CRISPR-Cas9. Red triangles indicated the cutting sites; **C.** The expression of *MYB* in HT-55 cells with CRISPR-Cas9 based disruption on the identified motifs; **D.** Demonstration of the motifs to be deletion by site-specific mutagenesis for the Luciferase assay. Red sequences indicated the targets of deletion; **E.** Western blotting showing the decrease in HNF4A protein levels upon *HNF4A* knockdown; **F.** The expression of *HNF4G* and *MYB* upon *HNF4G* knockdown. NC, non-targeted control. N = 3. **G.** The expression of *EBF1* and *MYB* upon *EBF1* knockdown. NC, non-targeted control. N = 3. *: P ⩽ 0.05; **: P ⩽ 0.01; ***: P ⩽ 0.001.

**Supplementary Figure 7. The genome-wide identification of MYB binding sites in HT-55 and SNU-719 cells**

**A.** DNA binding motifs identified using HOMER derived from MYB ChIP-seq in HT-55 and SNU-719 cells; **B.** Venn diagram showing the overlap of genes with MYB binding in HT-55 and SNU-719 cells; **C.** ChIP-qPCR analysis showing the enrichment of MYB at the enhancer e4 in HT-55 and SNU-719, GP2d and CCK-81 cells. NCR: Negative control region. N = 3; **D.** Luciferase reporter assay measuring the activity of the enhancer e4 upon *MYB* knockdown in HT-55 cells. N = 3; **E.** The MYB ChIP-seq tracks of MYB targeted cancer-related genes in HT-55 and SNU-719 cells. *: P ⩽ 0.05; **: P ⩽ 0.01; ***: P ⩽ 0.001.

**Supplementary Figure 8. The discovery of MYB targeted genes in HT-55 cells**

**A.** Volcano plot showing the differentially expressed genes in HT-55 cells upon *MYB* knockdown. Red:

upregulated genes with *MYB* knockdown. Blue: downregulated genes with *MYB* knockdown. Grey: unchanged genes; **B.** Venn diagram showing the genes which were bound and regulated by MYB; **C.** GSEA indicating that genes associated to Notch Signaling were downregulated upon *MYB* knockdown. NES, normalized enrichment score.

**Supplementary Figure 9. The repression of MYB expression in vivo prevented the development of gastrointestinal adenocarcinoma**

**A.** The expression of *MYB* upon *MYB* knockdown in HT-55 and SNU-719 cells; **B-C.** Western blotting showing the decreased MYB protein upon *MYB* knockdown **(B)** or CRSPRi targeting the enhancer e4 **(C)** in HT-55 and SNU-719 cells. The ACTB protein served as a loading control; **D.** RT-qPCR demonstrating the changes of Notch signaling related genes upon *MYB* knockdown. **E-G.** The growth of the xenografts based on HT-55 cells were significantly inhibited upon *MYB* knockdown. H. Immunohistochemical staining of Ki-67 in xenograft samples upon *MYB* knockdown in HT-55 cells. Scale bar = 250 μm. *: $P \leqslant 0.05$; **: $P \leqslant 0.01$; ***: $P \leqslant 0.001$.

### Supplementary figure 1.

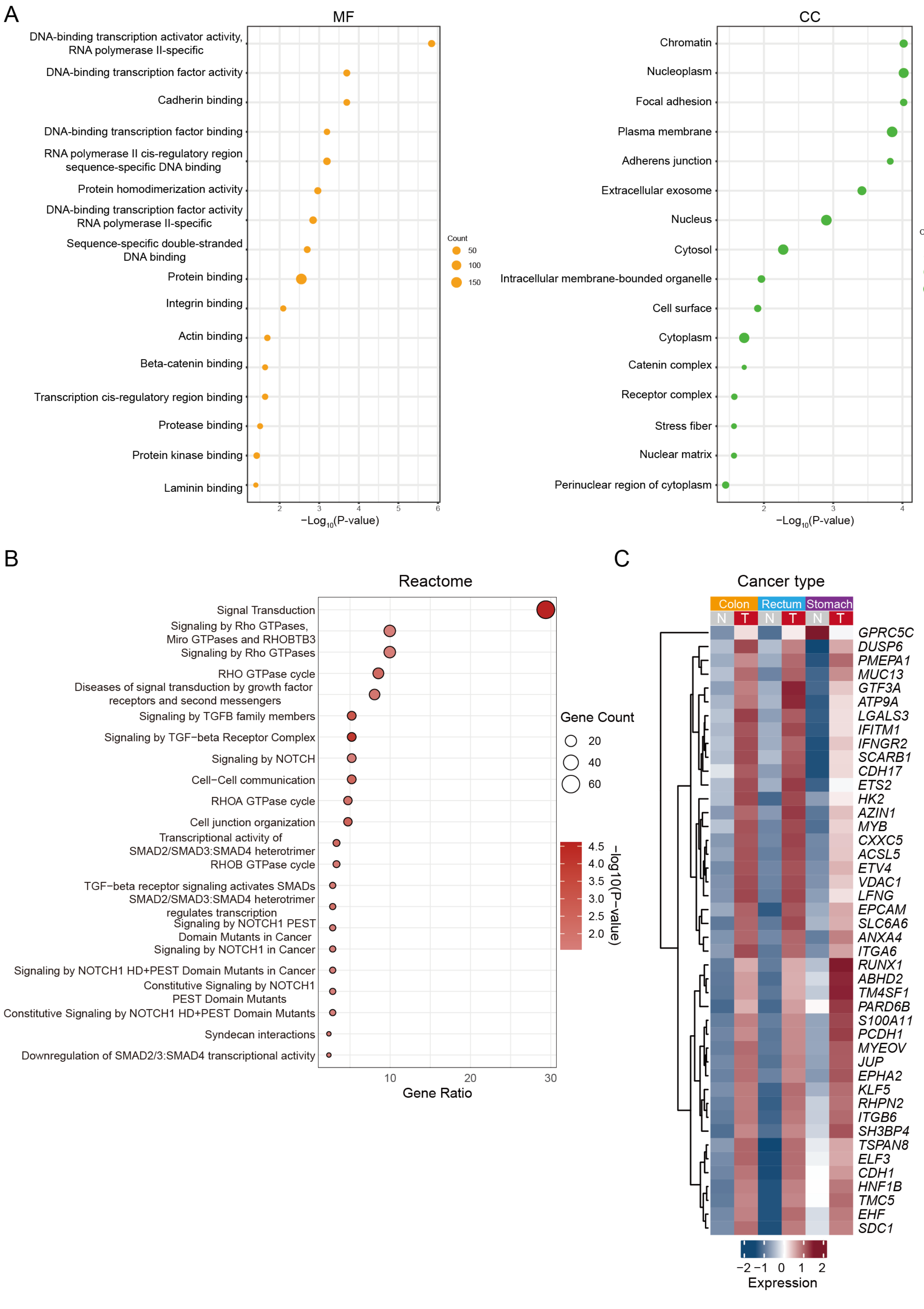

**Supplementary figure 2.**

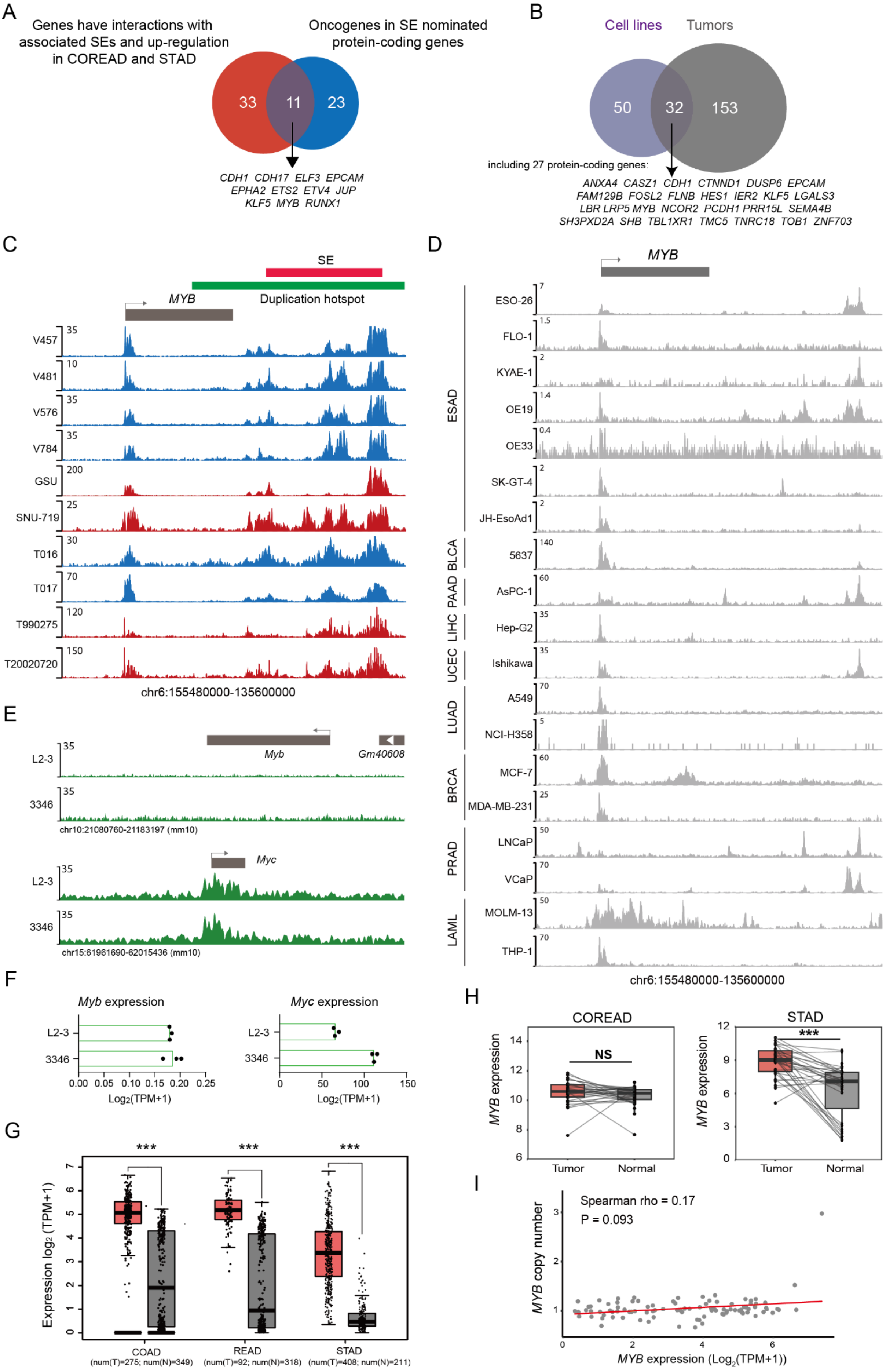

**Supplementary figure 3.**

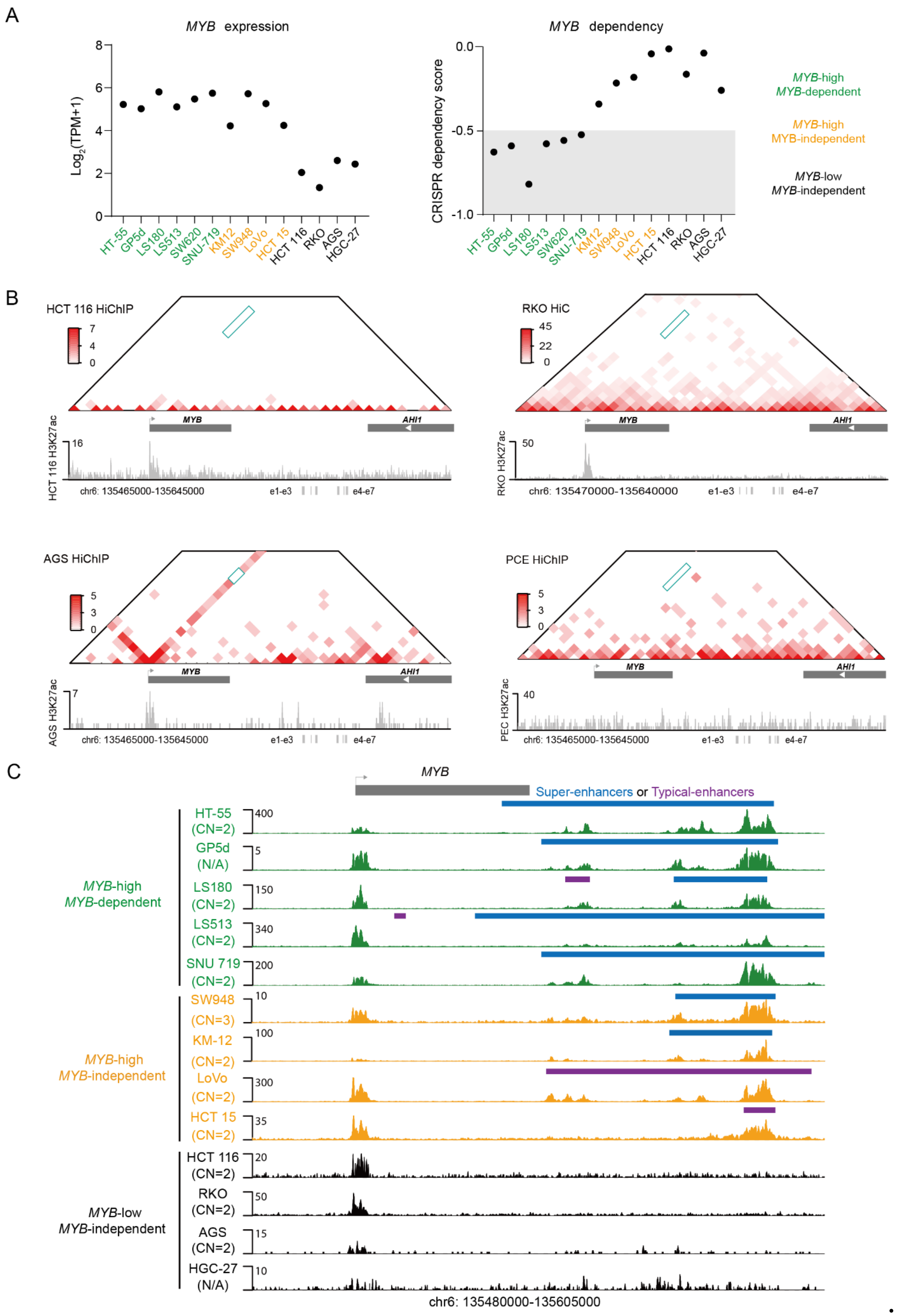

**Supplementary figure 4.**

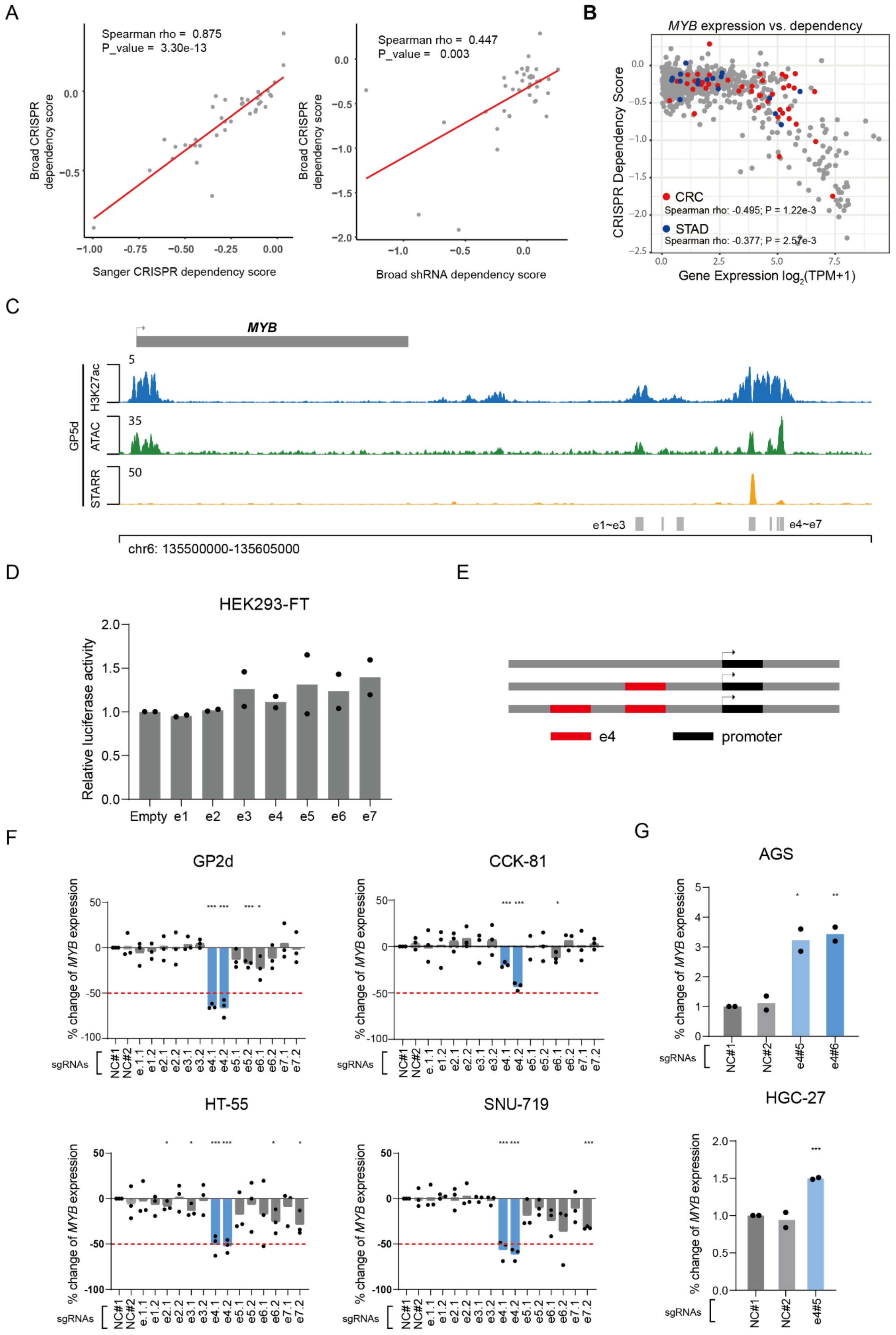

### Supplementary figure 5.

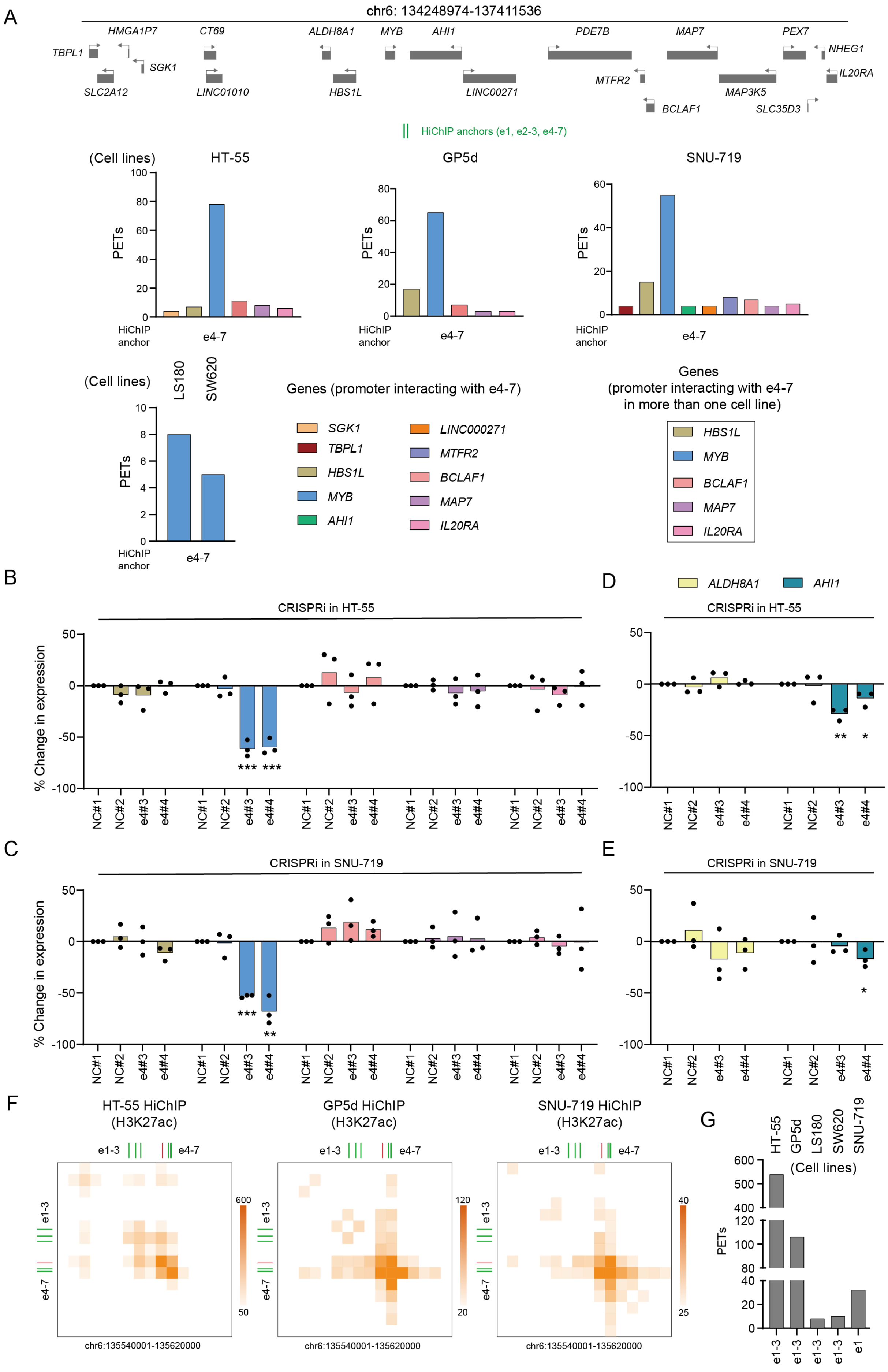

**Supplementary figure 6.**

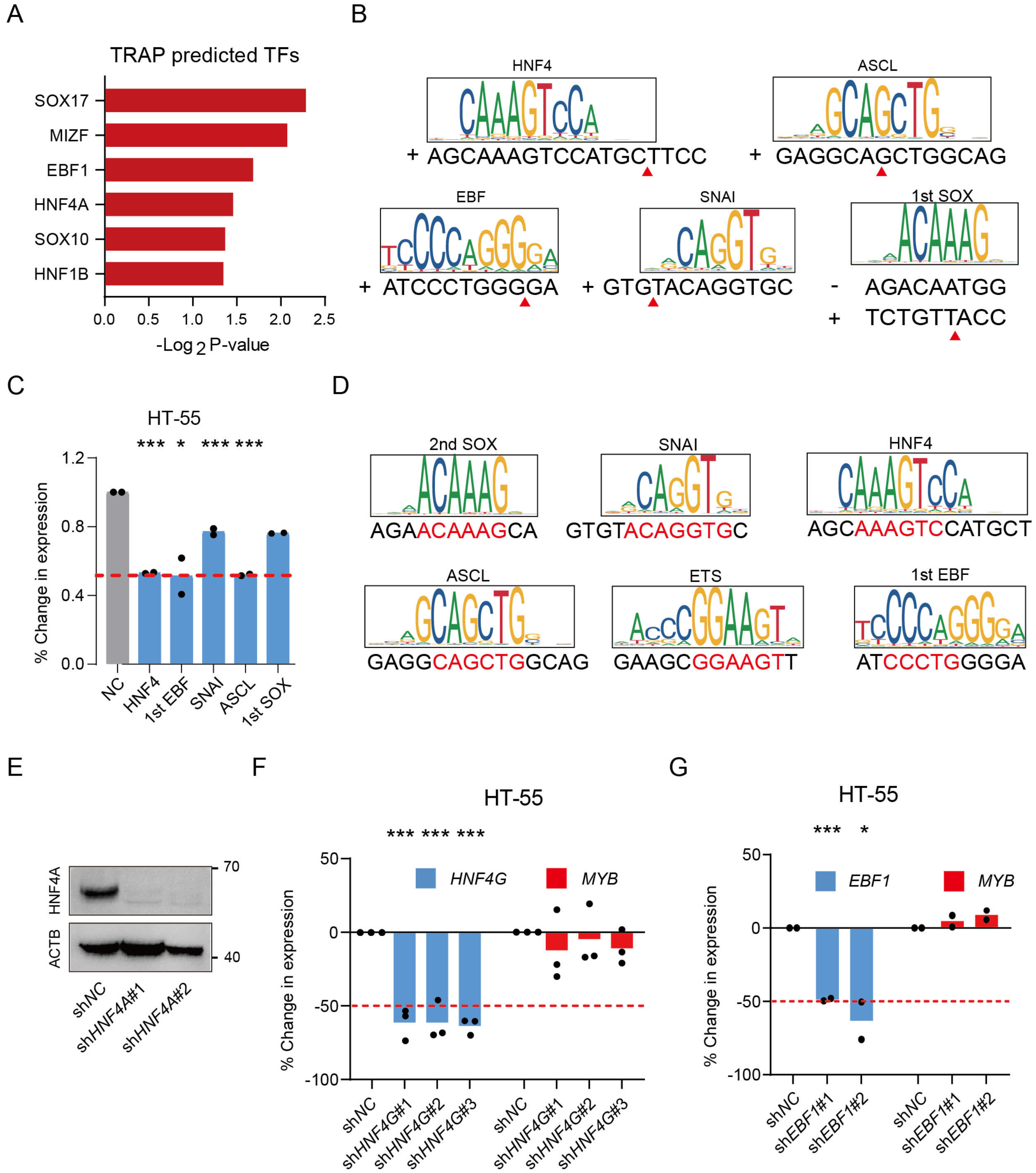

**Supplementary figure 7.**

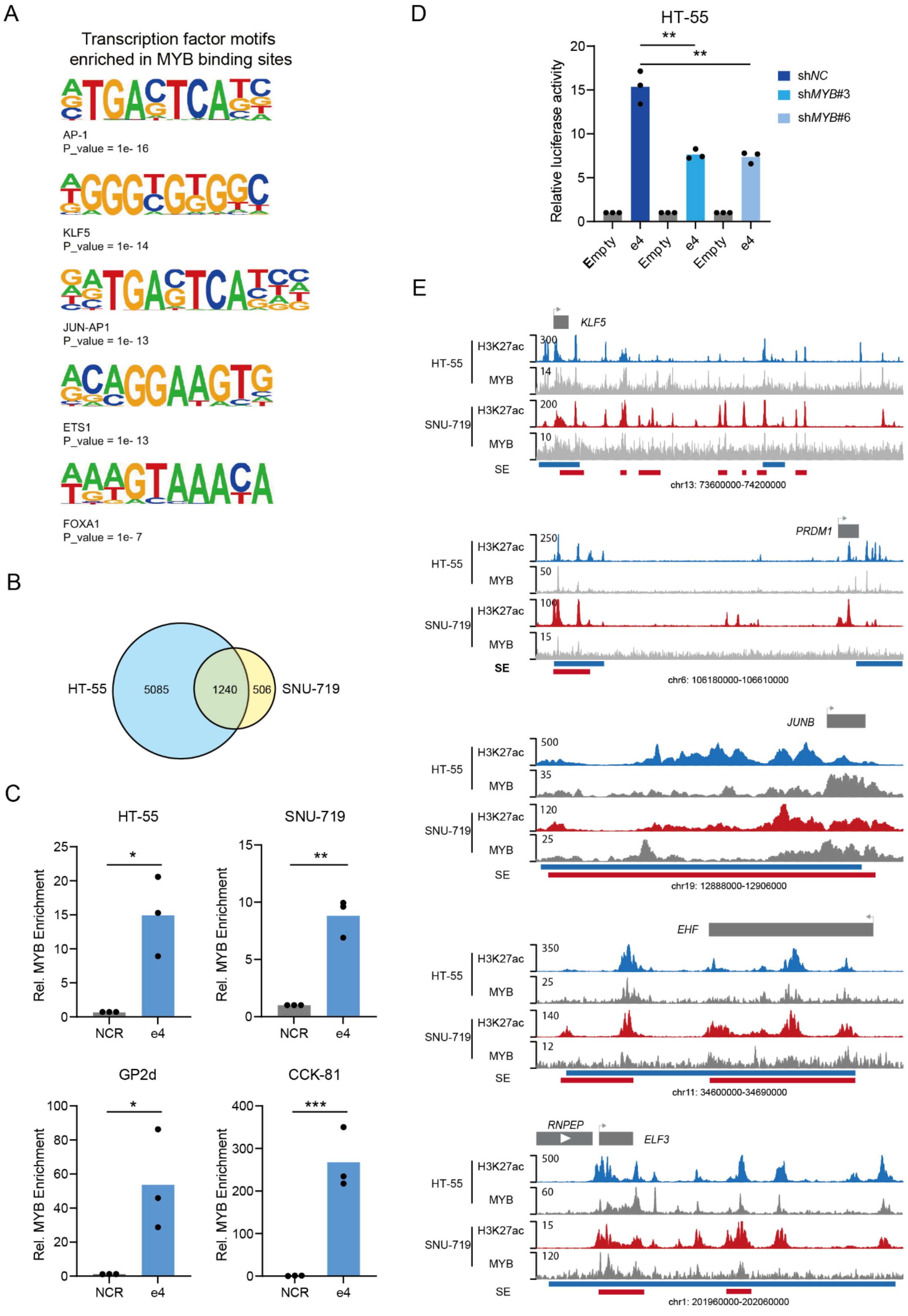

#### Supplementary figure 8.

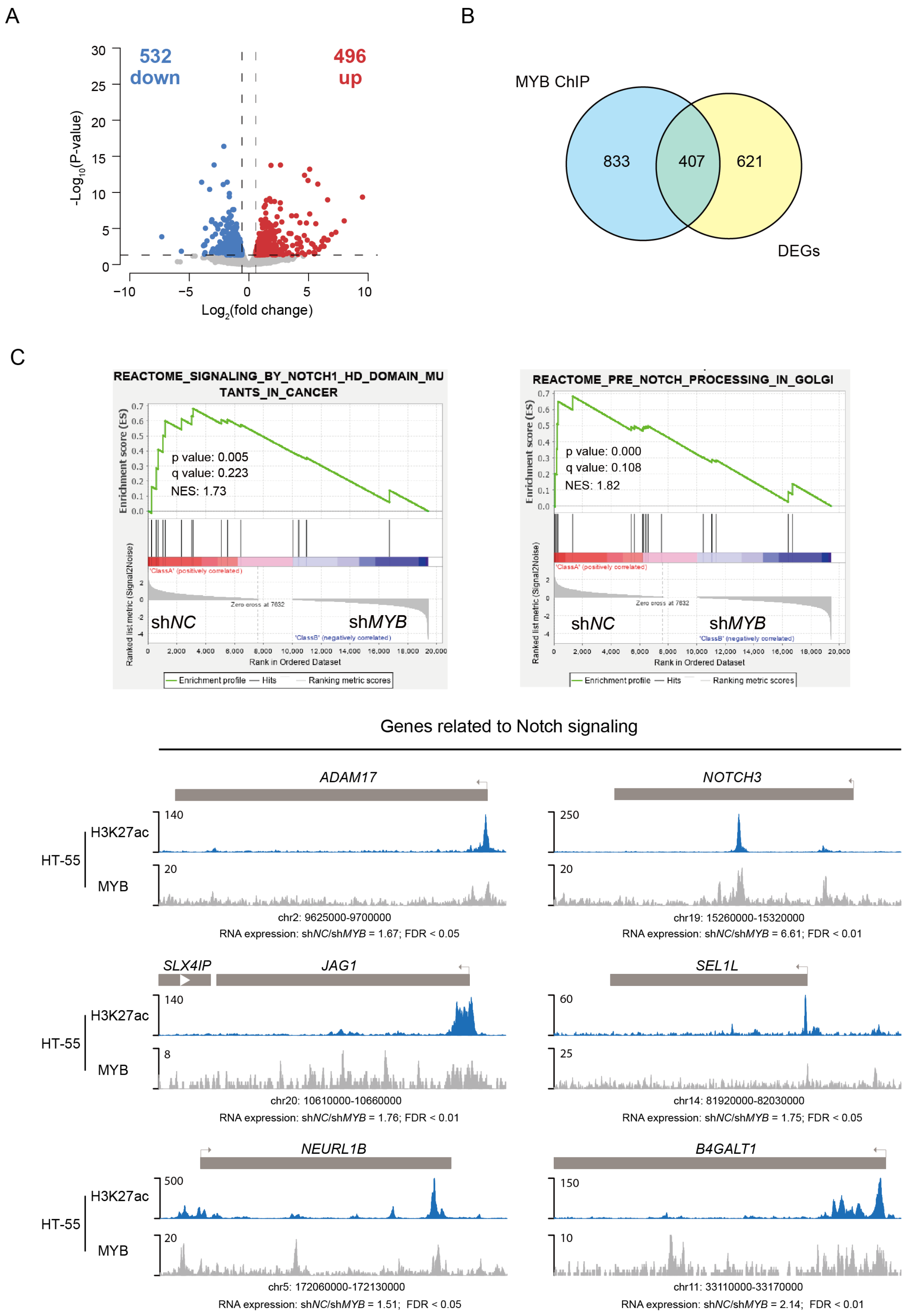

**Supplementary figure 8.**

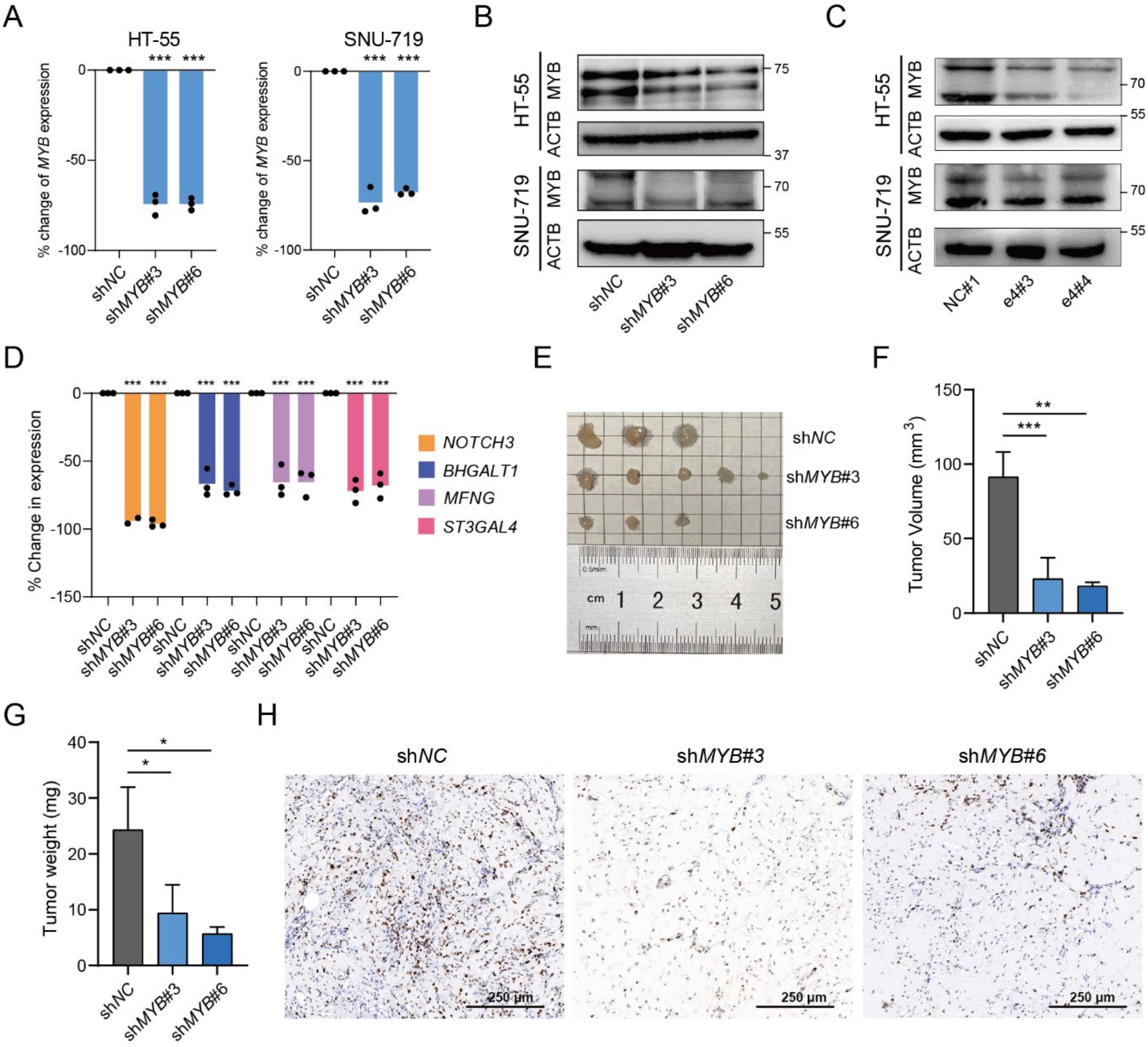
